# Uncovering the regulatory landscape of early human B-cell lymphopoiesis and its implications in the pathogenesis of B-cell acute lymphoblastic leukemia

**DOI:** 10.1101/2023.07.01.547234

**Authors:** Núria Planell, Xabier Martínez-de-Morentin, Daniel Mouzo, David Lara-Astiaso, Amaia Vilas-Zornoza, Patxi San Martín-Uriz, Diego Alignani, Bruno Paiva, Alberto Maillo, Aleksandra Kurowska, Nerea Berastegui, Paula Garcia-Olloqui, Arantxa Urdangarin, Peri Noori, Asier Ortega-Legarreta, Mikel Hernaez, Vincenzo Lagani, Narsis Kiani, Matthias Merkenschlager, Teresa Ezponda, José I. Martín-Subero, Ricardo N. Ramírez, Jesper Tegner, Felipe Prosper, David Gomez-Cabrero

**Affiliations:** Universidad de Navarra, Centro de Investigación Médica Aplicada (CIMA), Computational Biology Program, Instituto de Investigación Sanitaria de Navarra (IdiSNA), Pamplona, Spain; Translational Bioinformatics Unit, Navarrabiomed, Universidad Pública de Navarra (UPNA), Instituto de Investigación Sanitaria de Navarra (IdiSNA), Pamplona, Spain; Bioscience Program, Biological and Environmental Sciences and Engineering Division (BESE), King Abdullah University of Science and Technology KAUST, Thuwal 23955, Saudi Arabia; Hematology-Oncology Program, CIMA, Cancer Center Clínica Universidad de Navarra (CCUN), IdISNA, Pamplona, Spain. Centro de Investigación Biomédica en Red de Cáncer, CIBERONC; Flow Cytometry Core Facility, Centro de Investigación Médica Aplicada, Universidad de Navarra, CCUN, IDISNA, CIBERONC, Pamplona, Spain; Clínica Universidad de Navarra, Centro de Investigación Médica Aplicada, Universidad de Navarra, CCUN, IDISNA, CIBERONC, Pamplona, Spain; Translational Cardiology, Department of Medicine Solna, Karolinska Institute, Stockholm, Sweden; SDAIA-KAUST Center of Excellence in Data Science and Artificial Intelligence, Thuwal 23952, Saudi Arabia; Institute of Chemical Biology, Ilia State University, Tbilisi 0162, Georgia; Department of Oncology-Pathology, Center of Molecular Medicine, Karolinska Institutet, Stockholm, Sweden; Algorithmic Dynamics Lab, Center of Molecular Medicine, Karolinska Institutet, Stockholm, Sweden; MRC LMS, Institute of Clinical Sciences, Faculty of Medicine, Imperial College London, London, UK; Institut d’Investigacions Biomèdiques August Pi i Sunyer (IDIBAPS), University of Barcelona; Institució Catalana de Recerca i Estudis Avançats (ICREA), Barcelona, Spain; Department of Immunology, Harvard Medical School, Boston, MA 02115, USA; Computer, Electrical, and Mathematical Sciences and Engineering Division (CEMSE), King Abdullah University of Science and Technology KAUST, Thuwal 23955, Saudi Arabia; Unit of Computational Medicine, Department of Medicine, Center for Molecular Medicine, Karolinska Institutet, Karolinska University Hospital, L8:05, SE-171 76, Stockholm, Sweden; Science for Life Laboratory, Tomtebodavagen 23A, SE-17165, Solna, Sweden; Hematology and Cell Therapy Service, Cancer Center Clínica Universidad de Navarra (CCUN), IdISNA, Pamplona, Spain

## Abstract

Dysregulation of the early stages of B-cell lymphopoiesis, orchestrating the development of cellular immunity, may induce malignant transformations. Therefore, it is essential to characterize the gene regulatory network (GRN) driving B-cell lymphopoiesis in healthy individuals to uncover malignancy mechanisms. To this end, we generated a dataset that included paired human data for chromatin accessibility and gene expression in eight B-cell precursor stages, providing the first deep characterization of early B-cell lymphopoiesis, including the identification of regulatory elements and the reconstruction of the GRN. Using this data, we recapitulated well-known regulatory elements and revealed new regulons, such as ELK3, enriched in pro-B cells with a putative role in cell cycle progression. Moreover, a single-cell multi-omics analysis validated and enhanced the resolution of the regulatory landscape recovered by bulk data, revealing MYBL2 and ZNF367 as specific regulons of cycling cell states, and CEBPA associated with lymphoid multipotent progenitors (LMPPs). Importantly, this dataset enabled us to uncover B-cell acute lymphoblastic leukemia (B-ALL) triggers. We identified different cellular origins of malignant transformation depending on the B-ALL subtype, including the association of the ETV6-RUNX1 with pro-B cells and the increased expression of *ELK3* in this ALL subtype. Overall, our dataset provides the most comprehensive atlas to date of early human B-cell regulation (B-rex; https://translationalbio.shinyapps.io/brex/), facilitating further understanding of B-cell differentiation in health and disease.

## INTRODUCTION

B-cell lymphopoiesis gives rise to a diverse repertoire of peripheral B cells, antibody-producing cells capable of mediating protection against pathogens while remaining tolerant of self-antigens. Before migration to the periphery, B-cell development occurs in the bone marrow following a complex process governed by the sequential rearrangement of the immunoglobulin (Ig) heavy and light chain genes, and their expression and subsequent assembly of their encoded proteins into B-cell receptor (BCR)^1,2^. This highly ordered and controlled process is divided into several cell stages or B-cell subpopulations^3,4^. The transcriptional profile of each cell subpopulation emerges from an underlying gene regulatory network (GRN) in which a limited number of transcription factors (TFs) and cofactors regulate each other and their downstream target genes^5^. Several TFs orchestrating the B-cell development have been described^2,6^. Key lineage TFs, such as IKZF1 and PU.1, are implicated in the initial commitment towards the lymphoid lineage^7,8^. PAX5, EBF1, and E2A (TCF3) are regulators of B-cell lineage commitment^9–11^, while IRF4, IRF8, and FOXO1 are generally associated with Ig rearrangement^12,13^. Lastly, OCT2 and OBF1 are crucial TFs that drive the migration of immature B-cells to the periphery^14,15^. Most of this regulatory knowledge is derived from studies in mice^16–18^, and thus, it is unclear whether the mechanisms identified follow the same regulatory programs as those of humans^19^.

The integrated analysis of paired chromatin accessibility and gene expression profiling across various cell types offers a powerful approach to reconstructing GRNs and identifying cell-type specific regulons (TFs and their associated set of targets)^20,21^, a crucial approach to deciphering the regulation programs in humans. Such an analysis involves predicting cis-regulatory elements (CREs), identifying potential TFs, and linking the CREs to their candidate target genes. This strategy was followed to provide initial insights into the cis-regulation of mouse hematopoiesis^16^, capturing the key TFs committing the B-cell lineage, and in the study of mixed-phenotype acute leukemia, revealing RUNX1 as a putative regulator ^22^.

More recent advances in multi-omics single-cell approaches have allowed GRNs inference, applied to study human peripheral blood mononuclear cells (PBMCs)^20^ and plasma cell differentiation in the human tonsil^23^. Furthermore, a detailed characterization of B-cell subpopulations in mice and human lymphopoiesis has been obtained using single-cell technology approaches^3,4,24,25^. Despite significant progress, understanding of the regulatory mechanisms of this process remains incomplete^19,26,27^. In particular, current single-cell multi-omics experiments do not provide sufficient *depth* (detailed profiling) and *breadth* (enough individuals) to conduct detailed epigenetic profiling of the early stages of human B-cell differentiation.

To address this gap and to decipher the GRN of early B-cell differentiation in humans, we generated comprehensive chromatin accessibility and transcriptional profiles corresponding to the stepwise progression of B-cell precursors. Our study included eight cell subpopulations across 13 healthy donors. A detailed joint analysis of paired high-throughput data (bulk RNA-seq and ATAC-seq), allowed the generation of a comprehensive GRN of B cell differentiation, capturing cell-type specificity and dynamism of regulons, also further recapitulated using single-cell multi-omics. Interestingly, our data set recapitulated well-known regulatory elements and revealed novel core regulators of early B-cell differentiation, such as ELK3, associated with pro-B cell cycle progression. Finally, our in-depth characterization of normal human B-cell differentiation was ported to deconvolve B-cell malignancies; which served in deriving the transformation origin and putative regulators of some B-cell acute lymphoblastic leukemia (B-ALL) subtypes.

This detailed and extended portrait of regulatory elements has been made publicly available in a user-friendly web interface for browsing (B-rex; https://translationalbio.shinyapps.io/brex/), representing a rich playground to further unravel the regulatory mechanisms of B-cell differentiation in health and disease.

## RESULTS

### Profiling of early human B-cell differentiation stages by RNA-seq and ATAC-seq

To investigate the early B-cell differentiation in humans, we harvested bone marrow and peripheral blood from the same healthy individuals (n=13; mean age 21.4 [20-23]). Using fluorescence-activated cell sorting (FACS), we isolated hematopoietic stem cells (HSCs), common lymphoid progenitors (CLPs), pro-B cells, pre-B cells, and immature B cells from each bone marrow sample, and transitional B cells, and naïve B cells (CD5+ and CD5-) from peripheral blood specimens (**Figure 1A** and **Figure S1A**). All these populations were characterized using paired transcriptomic (RNA-seq) and chromatin accessibility (ATAC-seq) assays (**Figure S2A** and **Table S1**).

**Figure 1.**
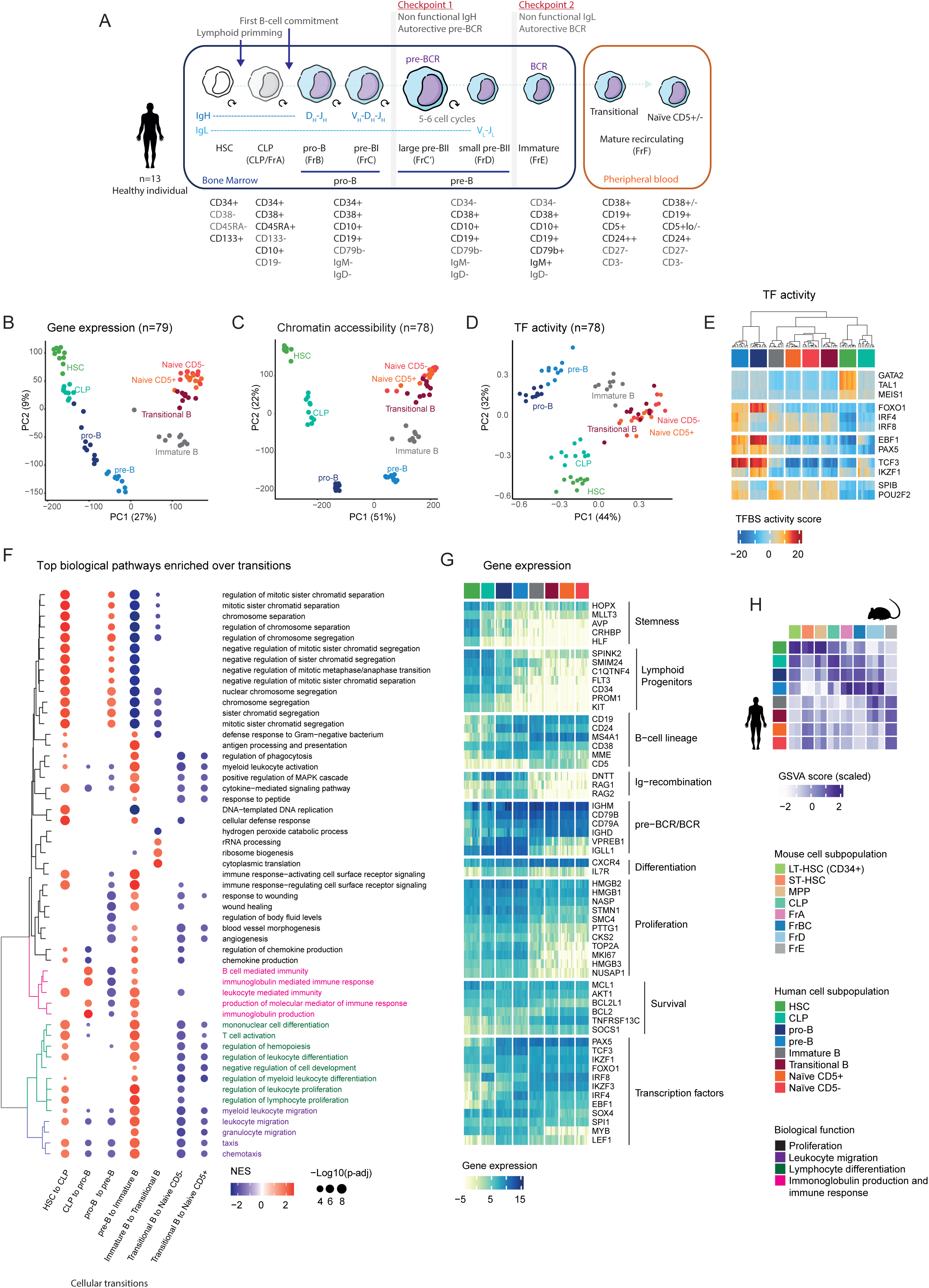
Characterization of early B-cell differentiation through chromatin accessibility and gene expression profiling. **(A)** Major B-cell precursor subpopulations from the human bone marrow and peripheral blood. The marker strategy followed to isolate 8 B-cell subpopulations from 13 healthy individuals is shown. Markers strongly expressed by the given B-cell subpopulation are depicted in black, whereas a lack of expression is shown in gray. **(B)** Principal Component Analysis (PCA) of gene expression (79 samples), **(C)** chromatin accessibility (78 samples), and **(D)** TFBS activity score (78 samples). Each dot represents a sample colored by cell subpopulation. **(E)** Heatmap representation of TFBS activity score for key B-cell regulators. Samples are shown by columns (n=78) and grouped by cell subpopulation, and TFs by rows (n=12). **(F)** Representation of top ten enriched GO terms derived from the transcriptional changes over B-cell transitions. The normalized enrichment score is shown in a red-blue color range for each term and the −log10 of the adjusted p-value as dot size. Biological functions are grouped into four main terms. **(G)** Heatmap representation of gene expression profile for 58 well-known genes associated with B-cell lymphopoiesis. Samples are shown by columns (n=79; grouped by cell subpopulation), and genes by rows (n=58; grouped by biological processes). Normalized gene expression levels are represented. **(H)** Heatmap representation of the gene set activity score (GSVA) computed for each human B-cell subpopulation signature (n=8; in rows) in each mouse sample (n=17; in columns). The GSVA score represented in the heatmap is scaled by column to highlight the human B-cell subpopulation with higher enrichment in each mouse B-cell subpopulation (in dark blue).

Principal component analysis of both chromatin accessibility and gene expression data resulted in a well-defined agreement in cell types across individuals recapitulating the dynamic process of B-cell differentiation (**Figure 1B, C**). We next evaluated transcription factor (TF) activity derived from the ATAC-seq data (TF binding site (TFBS); see Methods), which demonstrated a cell-type specific clustering of samples and recovered known TF-specific programs of B cell-differentiation (**Figure 1D, E**)^6^. Of all the transcriptional programs identified, pro-B and pre-B cells demonstrated the most divergent transcriptional program. This included immunoglobulin (Ig) rearrangement via FOXO1 and IRF4/8 and B-cell commitment with EBF1 and PAX5 TFs. Specific regulators of stemness such as GATA2, TAL1, and MEIS1 were observed in HSCs, the activation of lymphoid commitment was emphasized by the activity of TCF3 and Ikaros (IKZF1) in CLPs, and maturation in the late stages of differentiation by POU2F2 (OCT2) and SPIB (**Figure 1E** and **Figure S3** for the TFBS profile of all TFs).

To further assess the molecular processes captured by our dataset, we next examined biological pathways enriched in transcriptional changes over B-cell transitions (**Figure 1F** and **Table S2**) and the transcriptional and chromatin accessibility profiles of well-known markers of B-cell differentiation (**Figure 1G** and https://genome.ucsc.edu/s/TransBioLab/hg38_Brex for chromatin accessibility profile). We observed a significant enrichment of proliferation and differentiation pathways in CLPs maintained in pro-B and pre-B cells (**Figure 1F**). These pathways, in the early stages of B-cell differentiation, are mainly governed by the IL-7 receptor (IL7R) signaling, highly expressed in CLPs, pro-B, and pre-B cells, to later switch to the antagonistic signaling by the pre-B cell receptor (pre-BCR), expressed in pre-B cells (**Figure 1G**)^2,4^. Indeed, in pro-B to pre-B cell transition, we identified higher enrichment of proliferation pathways (**Figure 1F**), associated with pre-B cell expansion (large pre-B or FrC’) controlled by the convergence of IL7R and the pre-BCR signaling^28^. We also captured the fundamental process that organizes the B-lymphocyte development, the immunoglobulin (Ig) heavy-chain and light-chain rearrangement. We detected the enrichment of Ig-related pathways in the transition from CLPs to pro-B cells when the Ig rearrangement starts (**Figure 1F**). Accordingly, we observed the increased gene expression of *RAG1* and *RAG2* in pro-B and pre-B cells (**Figure 1G**). The successful Ig rearrangements derive in the B-cell receptor (BCR) expression by immature B cells and the migration to peripheral blood, which relates to the significant reduction of proliferation and the increased of migration pathways that we identified in pre-B to immature B cells transition (**Figure 1F**).

With a wealth of knowledge on B-cell differentiation stemming from studies described in mice, we next explored the conservation and translatability of our human B-cell transcriptional signatures (**Figure S4**) to those equivalents in mice (**Figure S5**)^29,30^. To achieve this, we measured a gene set activity score, a unitless measure where higher (lower) values are associated with a higher (lower) activity of a given gene set, derived from a Gene Set Variational Analysis (GSVA). We computed the activity score of each human B-cell subpopulation signature in each mouse sample. We observed a general concordance between species (**Figure 1H**), further supporting the sharing of transcriptional programs across human and mouse B-cell differentiation and development^31^.

Overall, we generated a rich picture of the gene expression and the transcriptional regulation via TF activity underlying the early B-cell differentiation in humans. This demonstrated well-preserved programs across individuals, matching the ones described in mice, showing the highly ordered processes subjacent to functional B-cell generation. We have prepared this atlas as a publicly available resource in a user-friendly web interface (B-rex; https://translationalbio.shinyapps.io/brex/).

### Defining the gene regulatory network and TF regulons driving early B-cell differentiation

Leveraging the collection of chromatin accessibility and gene expression provided the opportunity to study the underlying GRNs driving early B-cell differentiation in healthy humans. To this end, we defined a computational workflow to decipher the GRN from the RNA-seq and the ATAC-seq coordination. Briefly, it was generated through the inferred TFs accessible footprints mapping to CREs of differentially expressed genes across B-cell differentiation (**Figure S2B**). As a result, we obtained a single GRN for the entire B-cell differentiation involving 169 TFs, 7,310 genes, and 16,074 open chromatin regions (OCRs) (**TableS3** and **Figure S6A-C**).

Firstly, to explore the cell-type specificity of the regulons held in the GRN, we evaluated the Regulon Specificity Score (RSS)^20,32,33^, which measures the specificity of a regulon to a specific cell type based on the Jensen-Shannon divergence^34^ of target regions^20^ (a high RSS score indicates that the regulon is highly specific to the cell type). This analysis revealed high cell-type specificity of TF regulons (**Figure 2A**), capturing 5 main clusters of activator regulons according to TF expression and RSS. Several regulons were associated with HSCs including TAL1, MEIS1, MECOM, HLF, and GATA2 (**Figure 2A**, in cyan); others enriched over the differentiation process until reaching the pre-B cell stage, including GFI1, RUNX1, MYB (**Figure 2A**, in blue); few characteristics of pro-B cells such as ELK3 and BCL6B (**Figure 2A**, in green), the well-known SPI1, IRF4, TCF3, MEF2D, and EBF1 were mainly enriched in pre-B and immature cells (**Figure 2A**, in pink); and several associated to the late stages of differentiation, including PAX5, CEBPB, SPIB, etc. (**Figure 2A**, in black). Moreover, a repressive function was identified for 51 TF regulons, including PAX5, EBF1, and TCF3, according to previous knowledge (**Figure 2B**)^35^. To further understand the biological and molecular functions of the regulons, we performed a pathway enrichment analysis on the target genes within each regulon (**Figure 2C** and **Table S4**). We observed a significant enrichment of cell survival, proliferation, differentiation, and migration pathways for regulons associated with the first stages of differentiation (from HSCs to pre-B cells). Significant enrichment of pathways related to B lymphopoiesis and cell cycle was observed over the complete profile of regulons. Also, pathways related to cellular growth were enriched in those regulons highly expressed in pre-B cells, when large pre-B cells are generated, and bioenergetic pathways were enriched in regulons specifics from late stages of differentiation (from immature to naïve B cells).

**Figure 2.**
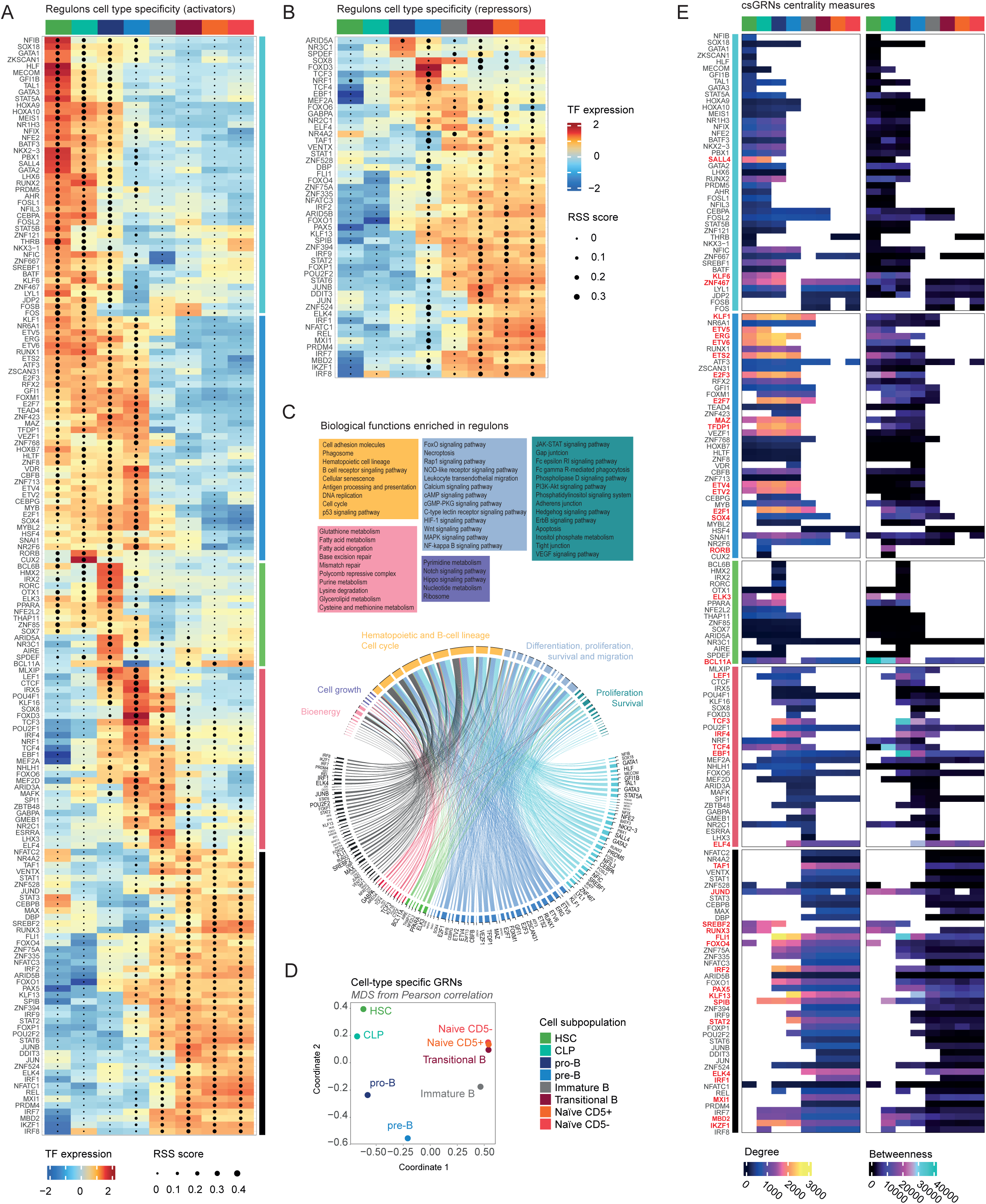
Regulatory landscape of TF regulons orchestrating early B-cell differentiation. **(A)** Heat map/dot-plot showing, for each TF regulon with an activator profile, the TF expression on a color scale and the Regulon Specificity Score (RSS) of target regions on a size scale. A high RSS score (high dot size) indicates that the regulon is highly specific to the corresponding cell type. TF regulons are shown by rows and B-cell subpopulations by columns. TF regulons are clustered into 5 main groups according to color and dot size scale (right side). **(B)** Heat map/dot-plot showing, for each TF regulon with a repressor profile, the TF expression on a color scale and the Regulon Specificity Score (RSS) of target regions on a size scale. A high RSS score (high dot size) indicates that the regulon is highly specific to the corresponding cell type. TF regulons are shown by rows and B-cell subpopulations by columns. **(C)** Circus plot summarizing the main biological pathways enriched for each regulon. On the bottom part are shown the TF regulons. TF regulons are clustered according to patterns identified in (A). On the top part are shown the enriched biological pathways, grouped into 5 blocks describing: bioenergetic processes (in pink), cell growth (in purple), hematopoietic and b-cell lineage and cell cycle (in yellow), differentiation, proliferation, survival and migration (in blue) and proliferation and survival (in turquoise). **(D)** Multidimensional Scaling representation from the Pearson correlation of cell subpopulation specific GRNs. Color-code by cell subpopulation. **(E)** Heatmap representation of the degree (left) and betweenness (right) centrality measures computed within each csGRN. TFs with the highest degree are shown in yellow, and those with the highest betweenness in cyan. Each column corresponds to a csGRN and each row to a TF regulon. TF regulons are clustered according to patterns identified in (A). The top 10 TF regulons of each csGRN based on degree and betweenness are highlighted in red.

We next sought to probe whether the cell-type specificity of TFs observed and their relative importance within the B cell-differentiation network could be further assessed using a rank-based analysis. We computed two measures of network centrality: (i) *degree*, which denotes the total number of edges that a TF has associated; and (ii) *betweenness*, which quantifies the influence of a given TF over the flow of information in the network (**Table S5;** see Methods). When we explored the top-10 ranked TFs by degree, we obtained several TFs from the ETS and Krüppel-like factor (KLF) family (EGR, FLI1, ETV5/6, ETS2, ELK4, and KLF1/13), implicated in a large number of cellular activities ^36,37^, TAF1, a key TF in the assembly of the preinitiation complex of the transcription process ^38^, and E2F3, a critical TF of cellular proliferation ^39^, all of them relevant for cellular survival. The top-10 ranked TFs based on node betweenness included critical TFs of B-cell differentiation as EBF1, TCF3, IRF4, and BCL11A ^6,40^, and some ETS and Krüppel-like factor (KLF) family TFs related to B-cell lymphopoiesis, including ERG, KLF13, ELK3, and ELK4^41–43^.

To further explore the key TFs in each cell subpopulation, we derived a specific GRN for each cell subpopulation (referred to as a cell-type specific GRN (csGRN); **Figure S2B** and **Figure S6D**). First, to validate that the retrieved csGRNs could recapitulate the B-cell differentiation trajectory, we computed distances between the csGRNs and generated a 2-dimensional projection. This csGRN driven embedding finely recapitulated differentiation trajectories observed when using the gene expression or chromatin accessibility data alone (**Figure 2D**). Next, we explored the centrality and betweenness measures for each csGRNs (**Figure 2E**). We observed how the csGRNs generally mirrored the regulon specificity shown in **Figure 2A**, as those TFs with low expression or chromatin accessibility in a specific cell state were excluded from the corresponding csGRN. Within the TFs with the highest values of degree and betweenness, we recovered the primary TFs implicated in B-cell commitment, differentiation, maintenance, or Ig recombination, such as TCF3, BCL11A, IRF8, PAX5, REL, IRF4, EBF1, POU2F2, FOXO1, SPB1, and LEF1 ^17,44–48^ (**Figure 2E**, in red).

Focusing on the regulons associated with pro-B and pre-B cells, subpopulations covering the lineage commitment to mature B-cells, we identified TCF3 as a key regulator of pre-B cells, together with EBF1, IRF4, and CTCF (cluster pink; **Figure 2A, C, E**). ELK3, PPARA, and BCL11A appeared as putative regulators of pro-B cells, controlling a large number of biological functions (cluster green; **Figure 2C**) and pointing to an important role in the transition from pro-B to pre-B stage (**Figure 2A, E**). The essential role of BCL11A in lymphoid development has already been reported ^40^, while the implications of PPARA and ELK3 in B-cells remain unknown. To note, the ETS-factor ELK3 has been associated with pro-B cell expansion in mice ^49^, and recently defined as a regulon in large and small B cells (FrC’/D - pre-B cells) in the Bursa of Fabricius of young broilers ^50^.

Taken together, we defined a multi-omics strategy able to recover the complex GRN of the entire B-cell differentiation process and the subjacent csGRNs, providing the dynamics of the regulons, recapitulating well-known regulatory elements, and uncovering new regulons with a putative role in B-cell lymphopoiesis, such as ELK3.

### Characterization of B-cell differentiation regulons revealed ELK3 as a cell cycle and differentiation regulator in pro-B cells

To further evaluate the value of this novel atlas in exploring the role of B-cell differentiation regulators, we paradigmatically first explored the well-known EBF1. Once we showed both the validity and the added value in EBF1, we then investigated the novel candidate regulator ELK3.

We identified eighty-five TFs as potential regulators of EBF1 (**Figure S7A**). Exploring those EBF1-regulators with the highest correlation with EBF1 gene expression, we found PAX5, TCF3, FOXO1, SPIB (Spi-1/PU.1 Related), RUNX1 and CEBPA, all of them well-known TFs controlling EBF1^51^. Next, exploring the EBF1-regulon, we identified 499 genes positively regulated, and 467 genes negatively regulated by EBF1. Within positively regulated genes, 18 were TFs, such as the already reported PAX5, TCF3, and FOXO1^52^ (**Figure S7B**). To unveil the biological functions of the EBF1-regulon, we performed a pathway enrichment analysis of the activated and repressed genes. This approach revealed a significant association of the EBF1-regulon-activated genes with B-cell activation, proliferation, and differentiation pathways, and the EBF1-regulon-repressed genes with metabolic and homeostatic processes (**Table S6**). Next, the pathway analysis of the EBF1-regulon for each csGRN resolved pathways significantly enriched in the pro-B, pre-B, immature, transitional, and naïve B cells csGRNs (**Figure S7C, D** and **Table S7**). Specifically, it highlighted the involvement of EBF1-regulon-activated genes in regulating cellular division in pro-B and pre-B cells. Moreover, it showed a role in regulating B-cell differentiation and activation through BCR signaling, starting from pre-B cells. Also, the EBF1-regulon-repressed genes showed specific metabolic regulation in pro-B and pre-B cells. In summary, this paradigmatic analysis of EBF1 manifested the power of our data for studying TF activity and regulatory control.

After confirming that our analysis recapitulates known characteristics of the well-established TF EBF1, we proceed to evaluate the potential role of ELK3 in human B-cell differentiation. ELK3 showed high gene expression levels and TF activity in HSCs, CLPs, and pro-B cells (**Figure 3A, B**), and emerged as a regulon within pro-B cells from our GRN analysis (**Figure 2**). We identified 50 TFs as potential regulators of ELK3, which included ERG, STAT5A/B, E2F3, ETS2, and ETV6, as activators, and SPIB (Spi-1/PU.1 Related) and POU2F2 as repressors (**Figure 3C**). Exploring the ELK3-regulon, we identified 1,339 genes positively regulated, and 1,294 genes negatively regulated by ELK3. Within regulated genes, 52 being TFs and including key regulators of B-cell differentiation such as PAX5, IKZF1, BCL11A, MEF2A, RUNX1, STAT5A, SPI1, and SPIB (**Figure 3D**)^53^. Next, to unveil the biological functions of the ELK3-regulon, we performed a pathway enrichment analysis of the activated and repressed genes. This approach revealed a significant association of the ELK3-regulon-activated genes with the cell cycle regulation, including the enrichment of several pathways related to DNA replication, unsaturated fatty acid metabolism, and MAPK and ERK cascade (**Table S8**). On the other hand, the ELK3-regulon-repressed genes showed a significant enrichment of several B-cell activation, proliferation, and differentiation pathways (**Table S9**). Importantly, when we assessed the pathway analysis of the ELK3-regulon for each csGRN resolved pathways there was a significant enrichment in the HSCs and pro-B cells. Specifically, it highlighted the involvement of ELK3-regulon-activated genes in regulating metabolic and cellular processes by the enrichment of fatty acid metabolic process, PI3K/PKB (Akt) signal transduction regulation, and MAPK cascade in both HSCs and pro-B cells. A significant enrichment of cell cycle pathways was observed in pro-B cells (**Figure 3E** and **Table S10**). The ELK3-regulon-repressed genes showed specific enrichment in pro-B cells, associated with several B-cell activation, proliferation, and differentiation pathways (**Figure 3F** and **Table S10**). Overall, our findings identify ELK3 as a previously underappreciated transcription factor in B-cell lymphopoiesis, which may play a crucial role in regulating the cell cycle and advancing B-cell differentiation during the pro-B cell stage.

**Figure 3.**
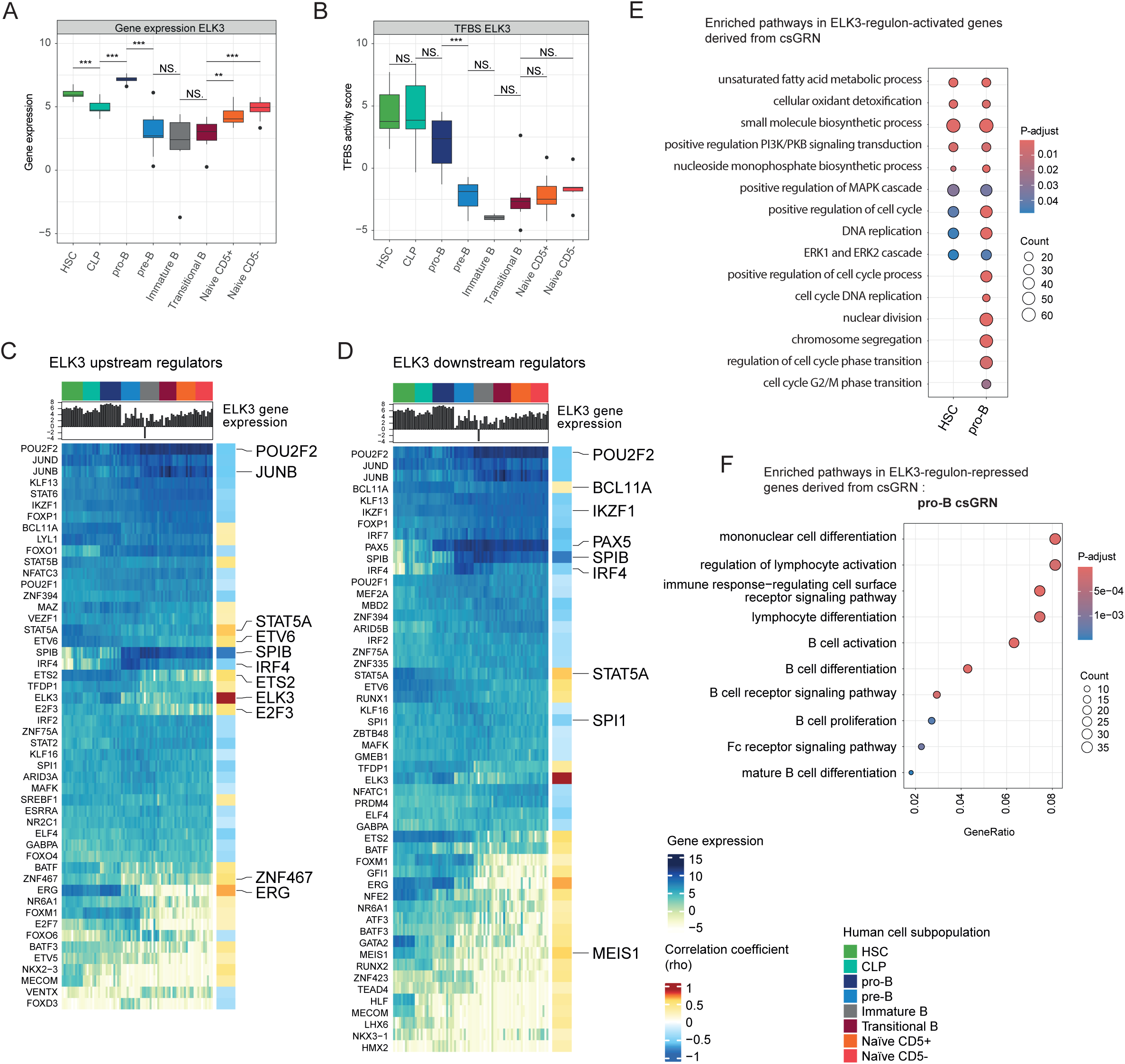
ELK3 regulon. **(A)** Box plot showing the distribution of ELK3 gene expression and **(B)** ELK3 TFBS activity score by cell subpopulation. Significant gene expression changes between cell subpopulation transitions assessed by the Wilcoxon test are shown. Significant changes were defined as a p-value < 0.05. (NS. = No Significant, “*” = 0.05, “**” = 0.01, “***” = 0.001). **(C)** Heatmap representation of gene expression for TF regulators of ELK3. Samples are shown by columns (grouped by cell subpopulation) and TFs by rows. Normalized gene expression levels are represented. On the right side rho values of Spearman correlation for each TF vs ELK3 are shown. On the top, a barplot representation of the ELK3 gene expression is shown. **(D)** Heatmap representation of gene expression for TF regulated by ELK3. Samples are shown by columns (grouped by cell subpopulation) and TFs by rows. Normalized gene expression levels are represented. On the right side rho values of Spearman correlation for each TF vs ELK3 are shown. On the top, a bar plot representation of the ELK3 gene expression is shown. **(E)** Representation of the main enriched GO terms derived from the pathway analysis of the ELK3-regulon-activated genes for each csGRN. The adjusted p-value is shown in a red-blue color range and the number of genes enriched within the term (counts) by dot size. **(F)** Representation of the main enriched GO terms derived from the pathway analysis of the ELK3-regulon-repressed genes for each csGRN. Significant terms only appeared in the assessment of csGRN derived from pro-B cells. For the pro-B csGRN enriched terms the adjusted p-value (red-blue color range), gene ratio, and counts (dot size) are shown.

### Multi-omics single-cell approach fine-tunes and validates bulk-derived regulons dynamics

Despite current limitations associated with single-cell sequencing technologies, including lower sequencing depth and capture efficiency, to increase the granularity of the early stages of B-cell differentiation and validate the observations from the bulk approach, we performed single-cell multi-omics analysis on FACS-sorted CD34+ hematopoietic progenitors isolated from 5 human samples (mean age 22.4 [21-24]) (**Figure 4A, Figure S1B,** and **Table S1**). For the single-cell RNA-seq (scRNA-seq) analysis, we recovered 28,891 cells representing undifferentiated cells and lymphoid lineage that clustered in 13 different cellular clusters. Those clusters were annotated as HSCs, multipotent progenitors (MPPs), lymphoid multipotent progenitors (LMPPs), CLPs, cycling pro-B cells, pro-B cells, cycling pre-BI cells, and pre-BI cells (**Figure 4B**), according to bulk-derived cell subpopulation profiles, known biomarkers, and cell cycle assessment (**Figure S8A-C**). For the paired single-cell ATAC-seq (scATAC-seq) analysis, we recovered 35,189 cells representing undifferentiated cells and lymphoid lineage, that showed a similar clustering and labeling as scRNA-seq (**Figure 4C**). Next, we performed a multi-omics analysis to identify the TF regulons from our single-cell data^20^. This analysis identified 62 regulons (**Table S11**), with 34 considered significant (21 activator (rho>0.7) and 13 repressor regulons (rho<-0.70); **Figure 4D** and **Figure S8D**). Most of the activator regulons identified matched with our bulk approach (17/21), including HLF, MECOM, MEIS1, NFE2, PBX1, RUNX1/2, CEBPA, THRB, LYL1, FOS, FOSB, MYBL2, LEF1, TCF4, EBF1, and PAX5. Interestingly, while we had identified MYBL2 regulon associated with pro-B and pre-B cells under bulk analysis (**Figure 2A**), here, we observed MYBL2 regulon highly specific to pro-B and pre-BI cycling cells (**Figure 4D, E**). Also, a finer regulon cell type specificity was identified for CEBPA, enriched in HSCs and CLPs under bulk approach (**Figure 2A**), but mainly associated with LMPPs in single-cell (**Figure 4D, E**). Within the new activator regulons appeared ZNF367, specific to cycling cells, and PLAGL1, NFIA, and HMGA2, mainly associated with HSCs, MPP, and LMPPs. In the case of repressor regulators, only PAX5 and EBF1 were identified by both approaches. However, the remaining significant repressor regulons showed poor coordination between TF expression and regulon accessibility (**Figure 4D**). Generally, this approach validated and enhanced the resolution of the regulatory landscape recovered by bulk and, although ELK3 regulon was not retrieved in our single-cell data, which may be a consequence of the CD34+ cells restricted approach, confirmed the specific increase of *ELK3* gene expression and chromatin accessibility in pro-B cells (**Figure 4F**).

**Figure 4.**
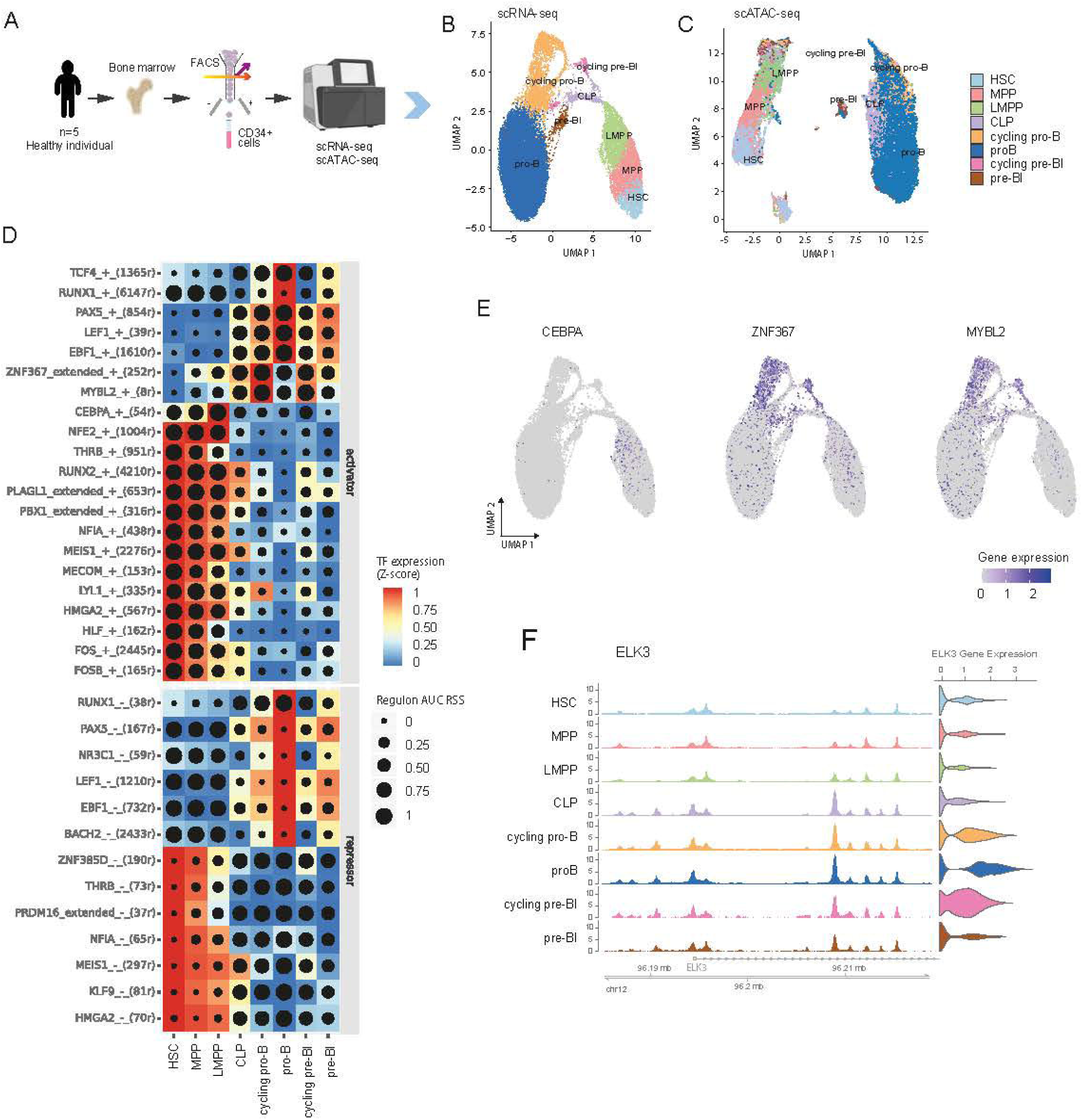
Regulation of early B-cell differentiation under multi-omics single-cell approach. **(A)** Schematic representation of data generated. Bone marrow samples were collected from 5 healthy individuals and CD34+ cells were isolated by flow cytometry. Then, each sample underwent a multi-omic single-cell approach, including scRNA-seq and scATAC-seq profiling. **(B)** Single-cell RNA-seq Uniform Manifold Approximation and Projection (UMAP) dimensionality reduction of 28,891 human CD34+ cells drawing the B-cell lymphopoiesis based on the annotated subpopulations. **(C)** Single-cell ATAC-seq Uniform Manifold Approximation and Projection (UMAP) dimensionality reduction of 35,189 human CD34+ cells drawing the B-cell lymphopoiesis based on the annotated subpopulations. **(D)** Heat map/dot-plot showing, for each TF regulon, TF expression on a color scale and the Regulon Specificity Score (RSS) of target regions on a size scale. A high RSS score (high dot size) indicates that the regulon is highly specific to the corresponding cell type. TF regulons are shown by rows and B-cell subpopulations by columns. TF regulons are clustered into activators (top) and repressors (bottom). **(E)** Single-cell RNA-seq Uniform Manifold Approximation and Projection (UMAP) dimensionality reduction of 28,891 human CD34+ cells drawing the B-cell lymphopoiesis based on gene expression of CEBPA, ZNF367, and MYBL2 genes. **(F)** Genomic data visualization including promoter region and initial bases of ELK3 gene (chr12: 96,185,000 – 96,219,000) (left) and violin representation showing the distribution of ELK3 gene expression (right) grouped by B-cell subpopulation.

### Identification of relevant B-cell subpopulations and regulons within subgroups of B-cell acute lymphoblastic leukemia subtypes

Given the comprehensive nature of our human B-cell profiling, we next turned our attention to the translatability of our results within a pathologic context. We hypothesized that the accurate characterization of the B-cell differentiation process would provide insights into the dysregulation of precursor B-cell differentiation-related diseases, such as B-cell acute lymphoblastic leukemia (B-ALL). Specifically, we aimed to use the provided framework of normal B-cell differentiation to identify the possible origin and mechanisms of malignant transformation of B-ALL subtypes. We investigated the enrichment of the transcriptional signature of eight public B-ALL subtypes (**Figure 5A**, **Figure S9A, B,** and **Table S12**)^54^ in our transcriptional signatures of normal B-cell precursors (**Figure S4A, B**) using an over-representation analysis (ORA). We found MEF2D fusions and TCF3-PBX1 ALL mainly enriched in pre-B cells, ETV6-RUNX1-like, Hyperdiploidy, and BCR-ABL1/ph-like in pro-B cells, KMT2A fusions subtype in CLPs, and DUX4 and ZNF384 fusions in HSCs (**Figure 5B**). We repeated the analysis exploring the enrichment of the B-ALL subtypes in our csGRNs, obtaining similar results (**Figure S9C**). Moreover, we computed the gene-set activity score for the derived B-cell subpopulation signatures in each B-ALL sample (**Figure 5C**), obtaining similar results to those observed in the ORA (**Figure 5B**). Additionally, we explored a list of 711 binding sites defined by Wray et al. as the highest affinity binding sites for ETV6-RUNX1 fusion ^55^ in our chromatin accessibility data, confirming the enrichment of ETV6-RUNX1 in pro-B cells (**Figure 5D**).

**Figure 5.**
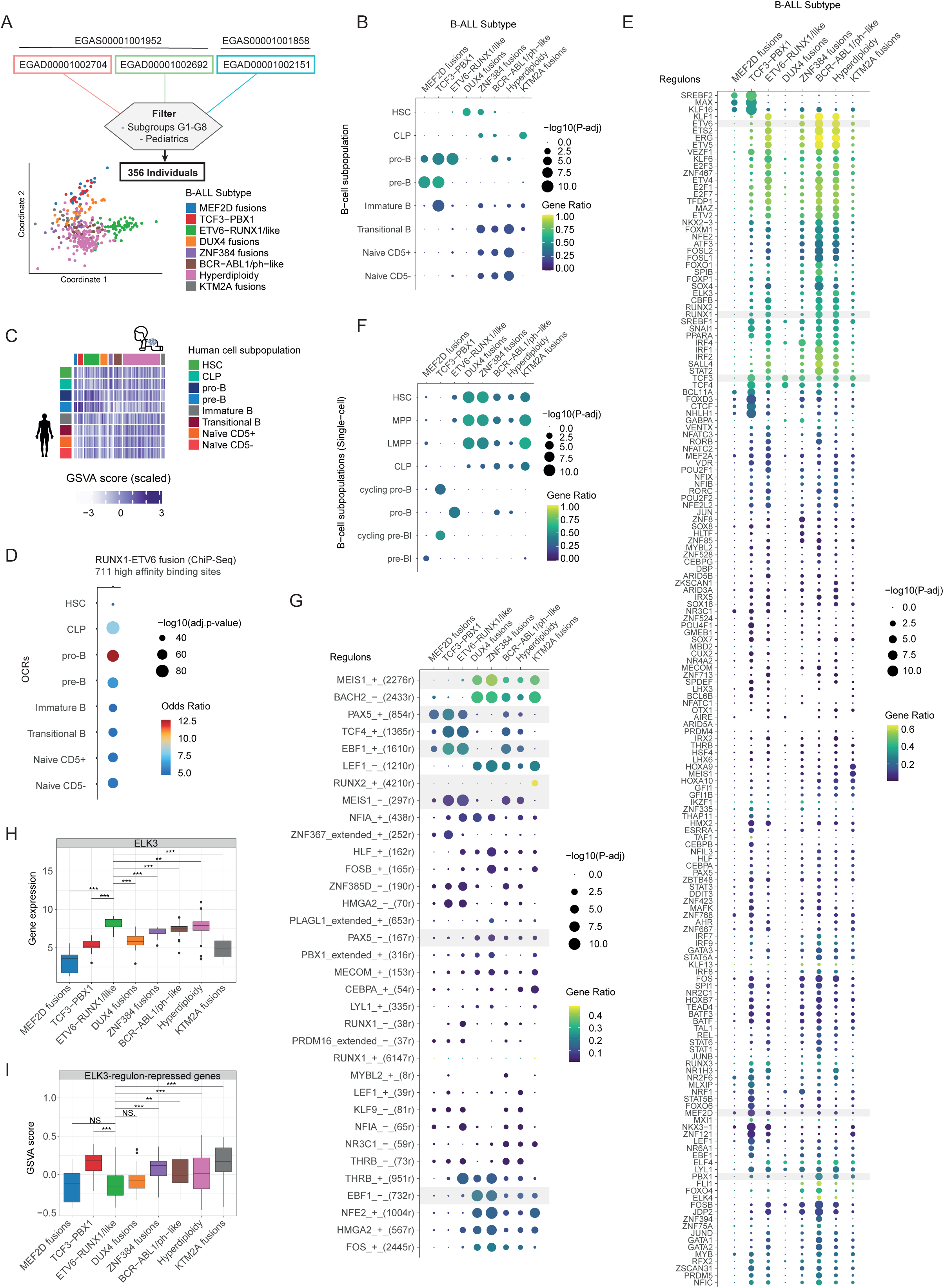
Uncovering B-ALL through normal B-cell differentiation regulatory landscape. **(A)** Schematic representation of B-ALL public data explored. **(B)** Over Representation Analysis (ORA) of B-ALL subtypes signatures over the B-cell subpopulations signatures. The dot size represents the −log10(adjusted p-value) and the color range the gene ratio. **(C)** Heatmap representation of the gene set activity score (GSVA) computed for each human B-cell subpopulation signature (n=8; in rows) in each B-ALL sample (n=356; in columns). The GSVA score represented in the heatmap is scaled by column to highlight the human B-cell subpopulation with higher enrichment in each B-ALL subtype (in dark blue). **(D)** ORA for RUNX1-ETV6 high affinity binding sites (n=711) over the consensus OCRs for each B-cell subpopulation. The dot size represents the −log10(adjusted p-value) and the color the odds ratio. **(E)** ORA of B-ALL subtypes signatures over the TF regulons defined by bulk approach (n=169). The dot size represents the −log10(adjusted p-value) and the color range the gene ratio. **(F)** ORA of B-ALL subtypes signatures over the single-cell B-cell subpopulations signatures. The dot size represents the −log10(adjusted p-value) and the color range the gene ratio. **(G)** ORA of B-ALL subtypes signatures over the TF regulons defined by single-cell approach (n=34). The dot size represents the −log10(adjusted p-value) and the color range the gene ratio. **(H)** Box plot showing the distribution of ELK3 gene expression by B-ALL subtypes. Significant gene expression changes between ETV6-RUNX1/like and the remaining B-ALL subtypes assessed by the Wilcoxon test are shown. Significant changes were defined as a p-value < 0.05. (NS. = No Significant, “*” = 0.05, “**” = 0.01, “***” = 0.001). **(I)** Box plot showing the distribution of the gene set activity score (GSVA) computed on the ELK3-regulon-repressed gene set by B-ALL subtypes. Significant changes between ETV6-RUNX1/like and the remaining B-ALL subtypes assessed by the Wilcoxon test are shown. Significant changes were defined as a p-value < 0.05. (NS. = No Significant, “*” = 0.05, “**” = 0.01, “***” = 0.001).

Next, aiming to decipher regulatory programs activated in each B-ALL subtype, we performed the TF regulons enriched in B-ALL subtypes (**Figure 5E**). Importantly, the TCF3-PBX1 subtype showed significant enrichment of TCF3 regulon and the ETV6-RUNX1-like subtype of ETV6 and RUNX1 regulons. Moreover, we identified similar enriched profiles for ETV6-RUNX1-like, BCR-ABL1/ph-like, and Hyperdiploidy B-ALL subtypes, reinforcing a common origin of the malignant transformation identified in **Figure 5B**. Also, TCF3-PBX1 and MEF2D fusions, both associated with pre-B cells, shared the top enriched regulons, including MAX, SREBF2, KLF16, and BCL11A. Few regulons were enriched for those B-ALL subtypes associated with the first stages of differentiation. Specifically, for DUX4 fusions where only TCF3/4 regulons were significantly enriched. For KTM2A fusions TCF3 but also HOXA9/10, MEIS1, ZNF121, and SREBF1 regulons showed a significant enrichment. In the case of ZNF384 fusions, a considerable number of regulons were enriched, being TCF3/4 and IRF4 those with the highest significance.

As some B-ALL subtypes showed enrichment with early stages of differentiation (like HSCs, CLPs, and pro-B cells), we next interrogated the enrichment of the B-ALL subtypes in our single-cell data from normal B-cell precursors (**Figure 5F**). This approach revealed the association of TCF3-PBX1 with cycling cells (both pro-B and pre-B cells), DUX4 and ZNF384 fusions with HSCs, MPPs, and LMPPs, and the KTM2A fusions strongly enriched in LMPPs. Within TF regulons, we identified similar profiles for DUX4, ZNF384, and KTM2A fusions, including MEIS1, BACH2, and LEF1 over the main significantly enriched regulons (**Figure 5G**). Moreover, significant enrichment of PAX5 was revealed inTCF3-PBX1, ETV6-RUNX1/like, and MEF2D fusions. Also, the enrichment of TCF4 and EBF1 in TCF3-PBX1 by bulk approach (**Figure 5E**) was confirmed by the single-cell approach (**Figure 5G**).

Following up on our earlier observations of ELK3 as a relevant TF in pro-B cells, which regulon has been reported overexpressed in ETV6-RUNX1-positive ALL^25^, we explored the gene expression of ELK3 and the activity score of ELK3-regulon over the B-ALL subtypes (**Figure 5H, I**). Interestingly, we observed an overexpression of ELK3 in ETV6-RUNX1/like ALL (**Figure 5H**), a subtype of B-ALL previously related to pro-B cells^25,56^. Moreover, although no changes were detected for the ELK3-regulon-activated genes (data not shown), a significant reduction of ELK3-regulon-repressed genes was observed in ETV6-RUNX1/like, DUX4, and MEF2D fusions (**Figure 5I**). Overall, these results demonstrate the utility of our comprehensive human atlas to obtain a deeper understanding of the molecular underpinnings of B-cell malignancies and sustain the hypothesis for the origins of B-ALL as a heterogeneous process that can arise from more primitive hematopoietic stem and progenitor cells ^24^.

## DISCUSSION

We addressed the challenge of profiling gene expression and chromatin accessibility during the early stages of B-cell lymphopoiesis in humans, to elucidate the underlying gene regulatory network (GRN). To this end, to our knowledge, we compiled the most comprehensive sample collection to date, encompassing bulk-derived multi-omics profiles of eight B-cell precursors from 13 healthy donors (EGAS00001007296), publicly available for browsing at https://translationalbio.shinyapps.io/brex/. This dataset, characterized by unprecedented granularity, provided the statistical power necessary for a detailed characterization of transcriptional and chromatin accessibility dynamics, allowing us to move beyond the transcription-based understanding of the gene regulatory landscape.

The definition of the B-cell precursors subpopulations has been well established in mice —the Hardy fractions—^29^, however, its translation to human surface markers remains controversial and an unequivocal assignment is missing^57,58^. Here, we defined a sorting strategy that allowed the recovery of the human cell fractions equivalent to HSCs, CLPs, pro-B cells (including FrB/C), pre-B cells (including FrC’/D), immature B cells (FrE), translational B cells (FrD), and naïve B cells (FrD) (**Figure 1A, H**), providing an unparalleled data set to robustly understand the system and complete what has been previously achieved in mice or mature stages of human B cells^16,18,59–62^.

Both gene expression and chromatin accessibility profiles delineate early B-cell differentiation and recovered well-known biomarkers, biological processes, and key TFs (**Figure 1**)^2,46^. Interestingly, chromatin accessibility, and more specifically intronic and distal intergenic regions, yield sharper distinctions between cell subpopulations than promoter regions or expression profiles (**Figure S10A, B** and **Figure S2A**), which is in line with previous observations in mouse immune system and human hematopoiesis^16,63^. This increased variability supports the crucial role of enhancer elements in the B-cell development regulation (**Figure S10C**).

Since gene expression and chromatin accessibility are not independent^64^, to recover the GRN of early B-cell differentiation, we defined a custom joint analysis inspired by a previous study in mice bulk data^65^ (**Figure S2B**). Although several assumptions underlie the computational approach and a fraction of false positive edges is expected, we consider such a network a powerful and very robust resource for hypothesis generation because (i) we used significant “open chromatin regions-gene” pairs derived from a large number of paired samples, (ii) we used footprinting, enabled by deep sequencing of ATAC-Seq (over 100 million reads per sample), to improve the estimation of active TF binding sites; (iii) the GRN recovers the already well-established regulators; and (iv) a specific GRN for each cell state can be derived permitting us to investigate further the cell-type specificity of the regulons (**Figure 2**). Moreover, we made use of public data to confirm that the defined cis-regulatory elements (CREs) were enriched by active histone marks^66^ and A-type compartments of chromatin conformation^67^ (**Figure S11A, B**). Also, using chromatin conformation data from naïve B cells^61^, we validated several CREs associated with characteristic genes from the late stages of differentiation, including MS4A1 (CD20), CD79A/B, PAX5, and IKZF1 (**Figure S11C**). However, Hi-C or promoter Hi-C experiments covering the early B-cell differentiation will be key in fine-tuning the defined GRN and allowing further interrogation of the cis-regulatory architecture leveraging deep learning models. Also, Chip-Seq experiments can help to validate the regulons identified.

The recovered GRN, not only captured the well-known regulators widely studied in mice but also shed light on understanding the B-cell differentiation process and the transition to malignant diseases. Interestingly, our GRN identified ELK3 as a new regulatory element of the early stages of B-cell differentiation (**Figure 2A, C, E**) implicated in the cell cycle progression mainly in pro-B cells (**Figure 3E**). This TF has recently been defined as a regulon in large and small B cells (FrC’/D - pre-B cells) in the Bursa of Fabricius of young broilers^50^, and it was associated with pro-B cell expansion in mice^49^ (**Figure 3H** and **Figure S9**). Moreover, ELK3 has been described as a novel oncogene reported increase in several carcinogenic processes including the ETV6-RUNX1-positive ALL, a kind of ALL that resembles the pro-B cell state and shows downregulation of the G2M checkpoint pathway ^45,56,68–71^. Importantly, ELK3 inhibition has been revealed as a potential candidate therapy for tumor growth and metastasis in mice and prostate cancer cells^72–74^. Although the putative implications of ELK3 in B-cell differentiation were unexpected, further understanding of the regulatory function of ELK3 merits evaluation.

To achieve a higher cellular resolution that is not feasible with sorting strategies, we transitioned to a single-cell approach, enabling the distinction between cycling and non-cycling cells. To note, although single-cell data covering the early human B-cell differentiation have already been published^4,22,25,75^, no one has yet used an RNA-ATAC multi-omics approach. This approach not only confirmed several regulons identified in the bulk analysis but also enhanced the specificity of their cellular contexts. However, when examining CD34+ cells using the single-cell method, the translation of our bulk results was somewhat limited, which might explain why the ELK3 regulon— showing the most significant changes during the pro-B to pre-B transition—was not fully recovered. It is important to highlight that our comprehensive profiling of bulk ATAC-Seq data played a crucial role in defining the open chromatin regions (OCRs) at the single-cell resolution, thereby adding robustness to the analysis.

The translational value of this novel comprehensive atlas was demonstrated in leveraging the novel healthy human B-cell differentiation insights to derive the putative cellular origin of the malignant transformation of several ALL subtypes. Our results support the heterogeneity of B-ALL origins and mirror the origin of malignant transformation recently described by Iacobucci et al.^24^. Additionally, the association of ZNF384 fusions with HSCs could explain the optimal treatment for this subtype of ALL patients, which currently is the allogenic-HSCT^76^, and the enrichment of hematopoietic stem cell features in TCF3-ZNF384-positive patients^77^. Interestingly, the single-cell data highlighted the association of ETV6–RUNX1 with no-cycling pro-B cells and the TCF3–PBX1 with cycling pro-B and pre-B cells (**Figure 5F**). Also, specific regulatory programs were identified for each B-ALL subtype, including the already known increase of HOXA9 and MEIS1 programs in KMT2A fusions^78^ and BACH2 in the transformation to BCR-ABL1/Ph1-like in mice^79^. Despite the interesting results, the unavailability of healthy samples in the B-ALL cohort made this approach challenging and limited the obtained results.

In summary, the generation of a complete and robust atlas of regulatory elements governing early human B-cell differentiation offers the ability to deeply interrogate the underlying gene regulatory networks and epigenetic components that finely distinguish healthy and disease cell states.

## MATERIAL & METHODS

### Human samples collection

Human bone marrow and peripheral blood samples from the same individuals were collected for 13 healthy donors (7 males and 6 females; median age of 21 [20-23]) (**Table S1**). Samples and data from donors included in the study were provided by the Biobank of the University of Navarra and were processed following standard operating procedures. The Clinical Research Ethics Committee of the Clinica Universidad de Navarra approved the study.

### FACS sorting

MNC cells were purified from whole blood and bone marrow samples using Ficoll-Paque PLUS (GE Healthcare. Illinois, USA). After that, eight immune cell populations, representing lymphoid hematopoietic differentiation cascade along the B cell lineage from adult human bone marrow precursors and peripheral blood, were obtained from each sample. These cells were purified following the sorting strategy shown in **Table S1** and **Figure S1A** in a BD FACSAria II (BD Biosciences) and then analyzed by bulk RNA-seq and bulk ATAC-seq.

### RNA-seq Data Generation

Bulk RNA-seq was performed following the MARS-Seq protocol adapted for bulk RNA-seq^80,81^ with minor modifications. Briefly, cells were sorted in 100 µl of Lysis/Binding Buffer (Ambion), vortexed, and stored at −80°C until further processing. Poly-A RNA was selected with Dynabeads Oligo (dT) (Ambion) and reverse-transcribed with AffinityScript Multiple Temperature Reverse Transcriptase (Agilent) using poly-dT oligos carrying a 7 nt-index. Up to 8 samples with similar overall RNA content were pooled and subjected to linear amplification by IVT using HiScribe T7 High Yield RNA Synthesis Kit (New England Biolabs). Next, RNA was fragmented into 250-350 nt fragments with RNA Fragmentation Reagents (Ambion) and dephosphorylated with FastAP (Thermo) following the manufactureŕs instructions. Next, partial Illumina adaptor sequences were ligated with T4 RNA Ligase 1 (New England Biolabs), followed by a second reverse transcription. Full Illumina adaptor sequences were added during final library amplification with KAPA HiFi DNA Polymerase (Kapa Biosystems). RNA-seq libraries quality controls consisted of quantification with Qubit 3.0 Fluorometer (Life Technologies) and size profile examination using Agilent’s 4200 TapeStation System. Libraries were sequenced in an Illumina NextSeq 500 in single-end at a sequence depth of 10 million reads per sample.

### RNA-seq Quantification and Normalization

Reads quality was assessed with FastQC software (Babraham Institute). Low quality, short reads, and TruSeq3-SE adapters were trimmed using Trimmomatic^82^ under the following parameters: ILLUMINACLIP:TruSeq3-SE:2:30:10 LEADING:3 TRAILING:3 SLIDINGWINDOW:4:15 MINLEN:30. Reads were then mapped to the December 2013 Homo sapiens high coverage assembly GRCh38 from the Genome Reference Consortium GRCh38 (hg38) using STAR [version 2.6.0a]^83^ and counted for each gene using htseq-count ^84^ with –m intersection-nonempty option and a GTF file from GRCh38 Ensembl annotation release 96, getting the information for a total of 58,884 genes. Genes with a minimum read count of 1 in all population replicates and an associated ensemble ID were retained (23,454 genes). A sample identified as an outlier has been removed (sample ID: D213-4). Counts were normalized under TMM normalization and voom transformation from edgeR r-package [v3.24.3] ^85^. Correlation between samples was computed using Pearson correlation (**Figure S10A**).

### RNA-seq Data analysis

Principal Component Analysis (PCA) was used for data exploration (**Figure 1B**). Differential expression analysis (DEA) was performed using limma [v3.38.3] and edgeR [v3.24.3] under two different approaches: 1) Identify biomarker genes for each cell subpopulation (all possible pair combinations). Genes with an adjusted p-value ≤ 0.01 (Benjamini-Hochberg) and absolute fold-change ≥ 1.5 at least in 4 contrasts were defined as significant (B-cell subpopulation biomarkers; **Figure S4**); 2) Identify differential expression genes (DEGs) between transition states (HSC vs. CLP, CLP vs. pro-B, pro-B vs. pre-B, etc.). Genes with an adjusted p-value ≤ 0.05 (Benjamini-Hochberg) and absolute fold-change ≥ 1.5 were considered significant. A clustering analysis was performed to identify the different expression patterns within the differentially expressed genes over transitions (second approach) using a model-base clustering based on parameterized finite Gaussian mixture models (mclust r-package [v5.4.5] under the following function and parameter: Mclust (modelName=”EII”)^86^) (**Figure S6A**). Gene set enrichment analysis of GO terms was performed using ClusterProfiler r-package [v4.6.0]^87^. The fold-change of DEGs at different transitions was given as input to the *compareCluster* function, and the top 10 enriched terms were plotted using an in-house *treeplot* function (**Figure 1F**).

### ATAC-seq Data Generation

Accessible chromatin mapping was performed using FAST-ATAC-seq^63^ with minor modifications. Briefly, 10,000 cells were sorted in 1X PBS, 0.05% BSA and pelleted by centrifugation at 500 rcf for 5 min at 4 °C with low acceleration and brake settings in a pre-cooled swinging-bucket rotor centrifuge. All the supernatant but 5 µL was removed. Next, 25 µL of transposase mix (15 µL of 2x TD buffer (Illumina); 1 µL of TDE1 (Illumina), 0.25 µL of 5% digitonin (Promega); 8.75 µL of nuclease-free water) were added to the cells, and the pellet was disrupted by pipetting. Transposition reactions were incubated at 37 °C for 30 min in an Eppendorf ThermoMixer with shaking at 450 rpm. Reactions were stopped at 4 °C for 5 min. To release tagmented DNA, samples were incubated at 40 °C for 30 min with 5 µL of clean-up buffer (900 mM NaCl (Sigma), 30 mM EDTA (Millipore), 2 µL of 5% SDS (Millipore) and 2 µL of Proteinase K (NEB)). DNA was purified using a 2X SPRI beads cleanup kit (Agencourt AMPure XP, Beckman Coulter). Two sequential PCRs were performed using KAPA HiFi DNA Polymerase and customized Nextera PCR indexing primers (IDT) ^88^. The conditions of the first PCR were: 5 min at 72 °C and 2 min at 98 °C followed by 9 cycles of 20 secs at 98 °C, 30 secs at 63 °C, and 1 min at 72 °C. Depending on the library concentration in this first PCR, a second PCR (2 min at 98 °C followed by 4 to 6 cycles of 20 secs at 98 °C, 30 secs at 63 °C, and 1 min at 72 °C) was performed aiming for a library concentration in the range of 2 to 10 ng/µL. PCR products were purified using a 2X SPRI beads cleanup. Libraries were quantified, and their size profiles were examined as described above. Sequencing was carried out in two batches, the first one in an Illumina NextSeq 500 using paired-end, dual-index sequencing (Rd1: 38 cycles; Rd2: 38 cycles; i7: 8 cycles; i5: 8 cycles) at a depth of 10 million reads per sample and second one in a HiSeq2500, v4 reagents using paired-end, dual-index sequencing (Rd1: 125 cycles; Rd2: 125 cycles; i7: 8 cycles; i5: 8 cycles) at a depth of 100 million reads per sample.

### ATAC-seq Quantification and Normalization

Reads quality was assessed with FastQC software (Babraham Institute). After trimming NexteraPE-PE adaptor sequences and low-quality reads and cutting the reads from the second batch to 38 bps using trimmomatic^82^ under the following parameters: CROP:38 ILLUMINACLIP:NexteraPE-PE.fa:2:30:10 LEADING:3 TRAILING:3 SLIDINGWINDOW:4:15 MINLEN:15. Reads were then mapped to hg38 reference genome (GRCh38 Ensembl annotation release 96) using BWA-MEM ^89^. Unmapped, unknown, random, ChrM mapping, duplicate reads, and reads matching to black list (the blacklist includes problematic regions of the genome^90^) were filtered out using samtools view -b -F 4 -F 1024 (samtools version 0.1.19), and Picard Tools (Picard MarkDuplicates, http://broadinstitute.github.io/picard). BAM files from re-sequenced samples were merged using samtools merge. Paired-end reads were used to determine those regions with significant reads coverage in contrast to the random background (regions known as peaks or Open Chromatin Regions; OCRs) using MACS2 (function *callpeak*) with the following parameters: -g hs -f BAMPE -q 0.01 --nomodel --shift -100 --extsize 200 -B –SPMR (https://github.com/taoliu/MACS) ^91^. A minimum OCR size of 20 bps was set as a threshold. To get the consensus OCRs for each cell subpopulation, we found the genome regions which are covered by at least n-1 of the sets of peaks (n being the total number of samples for a specific cell subpopulation), and then we merged the nearby peaks with a min gap width of 31. To get the consensus OCRs in the whole population, we get the regions in the genome where coverage is at least in 1 of the consensus OCRs from the different cell subpopulations and then merged the nearby peaks with a min gap width of 31, resulting in 105,530 ATAC-seq OCRs. These processes were done using the rtracklayer [v1.58.0] and GenomicRanges [v1.50.2] r-packages^92,93^.

BigWig files were generated for each BAM file using the *bamCoverage* function (--binSize 20 -- normalizeUsing RPKM --effectiveGenomeSize 2913022398) from deepTools [v3.3.1] ^94^. Then, a merged BigWig for each cell subpopulation was obtained through *bigWigMerge* from UCSC (http://hgdownload.cse.ucsc.edu/admin/exe/macOSX.x86_64/). Also, BigBed files were generated from consensus BED files of each cell subpopulation using *bedToBigBed* from UCSC. These data were deposited in a public custom track from UCSC (https://genome.ucsc.edu/s/TransBioLab/hg38_Brex).

Using the whole consensus OCRs, we compute the count matrix using the *multicov* function from bedtools [version 2.25.0] ^95^. Those OCRs without a minimum of 10 counts for at least one sample and 15 total counts were excluded (155 OCRs), retaining 105,359. Then, we normalize the data using the TMM method from edgeR r-package [v3.30.3] ^85^. One of the samples analyzed was identified as an outlier and excluded (sample ID: D215-5). Correlation between samples was computed using Pearson correlation (**Figure S10B**).

### ATAC-seq Data analysis

Principal Component Analysis (PCA) was used for data exploration (**Figure 1C**). OCRs were mapped to genomic locations (promoter, intronic, distal intergenic, etc.) using the ChIPseeker r-package [v1.18.0] ^96^ under the following arguments: Region Range of TSS as +/− 1000 bps, gene level, “TxDb.Hsapiens.UCSC.hg38.knownGene” TxDb object [v3.4.0] and “org.Hs.eg.db” annotation package [v3.7.0] (**Figure S2A**). The Gini index was computed for each OCR to measure “chromatin inequality” (**Figure S10B**). A Gini index of 0 represents perfect equality and similar chromatin accessibility in all cell subpopulations; while a Gini index of 1 represents perfect inequality, all the accessibility OCRs come from a specific cell subpopulation, and the others show no accessibility. To compute the Gini index, the maximum chromatin accessibility observed from the normalized count matrix was assigned for each cell subpopulation. Then, the *gini* function from REAT r-package [v3.0.2] was applied ^97^. Differential accessibility analysis (DAA) was performed to identify differential accessibility regions between transition states (HSC vs. CLP, CLP vs. pro-B, pro-B vs. pre-B, etc.) using a quasi-likelihood negative binomial generalized log-linear model (glmQLFit) from edgeR [v3.24.3] ^85,98^. OCRs with an adjusted p-value ≤ 0.01 (Benjamini-Hochberg) and absolute fold-change ≥ 2 were defined as significant. A clustering analysis was performed to identify the different accessibility patterns within the differentially accessible regions under a model-base clustering based on parameterized finite Gaussian mixture models (mclust r-package [v5.4.5] under the following function and parameter: Mclust(modelName=”EII”))^86^ (**Figure S6B**). To explore the accessibility of TFs, we computed the aggregated motif score (TFBS accessibility score) for each TF using chromVAR r-package [v1.5.0] ^99^ with the default parameters. For plotting the TFBS score was used (**Figure 1E** and **Figure S3**).

### Gene Set Variation Analysis (GSVA)

We performed the Gene Set Variation Analysis (GSVA) to compute the enrichment of a specific gene set within each single sample^100^. We applied it to explore the transcriptional proximity of the different B-cell states between humans and mice (**Figure 1H**). To that end, we used the defined biomarker gene sets for each B-cell subpopulation in humans (**Figure S4**) and computed the GSVA enrichment scores using the algorithm of Hänzelmann, Castelo, and Guinney (2013) by calling the gsvaParam() function from the GSVA R package^100^. We also used the GSVA to assess the transcriptional proximity of the different B-cell states and the B-ALL subtypes (**Figure 5C**).

### OCR Variance Component Analysis

We used a linear mixed model (LMM) approach for assessing the proportion of variance in gene expression explained by promoter and distal enhancer OCRs’ activity (**Figure S10C**). For each gene, we fit the following LMM regression:

Y = U_promoter + U_enhancers + Epsilon

where Y is a vector of zero-centered gene expression values, U_promoter and U_enhancer are random effects, and Epsilon explains noise unaccounted by the model. The limix python library was used for fitting the regression models (https://github.com/limix/limix) ^101^.

### OCRs association with target genes (cis-regulatory elements)

To determine the cis-regulatory elements (CREs) of a gene, we follow two approaches: 1) For promoter OCRs, we directly annotate OCRs to genes based on TSS distance (<1kb), and 2) For intronic or distal intergenic OCRs, we explored the association between chromatin accessibility and gene expression. First, we identified all intronic or distal intergenic OCRs for each expressed gene within a window of ±1Mb from the gene’s TSS. Second, the Spearman correlation was computed across all samples (59 samples; 8 cell subpopulations) for each gene-OCR pair (**Figure S2B**). Bonferroni correction was used to adjust for multiple testing. Gene-OCR pairs with adjusted p-value < 0.05 were considered significant.

### OCRs association with transcription factors (trans-regulatory elements)

To annotate OCRs to putative transcription factor binding motifs, we first identify the footprints within our OCRs. For each sample, we recognize the footprints using two algorithms, HINT^102^ and Wellington ^103^, using as input the corresponding sample BAM file and BED file from the corresponding cell subpopulation. For the footprints identified by HINT, we filtered out those footprints with less than 20 counts. Then, we get the union of the footprints identified for each sample through both strategies. Following, we compute the consensus footprint for each cell subpopulation following a similar strategy as for OCR consensus. To get the consensus footprint for each cell subpopulation, we found the genome regions covered by at least n-1 of the sets of regions (n being the total number of samples for a specific cell subpopulation). Finally, to get the consensus footprint in the whole population, we get the regions in the genome where coverage is at least in 1 of the consensus footprint from the different cell subpopulations; then, those with a width less than 6bps were removed, resulting in a total of 380,922 ATAC-seq footprints. These processes were done using the rtracklayer [v1.58.0] and GenomicRanges [v1.50.2] r-packages^92,93^. The identified footprints were located in 72,771 OCRs (69%).

Identified the footprints, we want to annotate the putative transcription factor binding motif to each footprint. To do that, we used the Vierstra database^104^. First, we generate a reduced version of the Vierstra database, getting those annotations falling within our OCRs regions using TABIX^105^. Then, we annotated at the archetype level (v1.0) to reduce motif redundancy and overlap, resulting in 30,258,213 region-motif annotations and 286 motif archetypes. The overlap between footprints and motif archetypes was explored, defining a minimum overlap of 90% of archetype width. Finally, footprints were linked to the original OCR to get the OCR-Archetype matrix, resulting in a matrix 105,530 x 282.

To translate archetypes to TFs, we define the following strategy. First, we compute the aggregated motif score (TFBS accessibility score) for each archetype using chromVAR r-package [v1.5.0]^99^ with the default parameters. Then, to identify the TFs most likely associated with an archetype, we compute, for each archetype, the Spearman correlation between the TFBS score of the archetype and the RNA expression of each TFs clustered in that archetype. Correction for multiple comparisons was done using the Benjamini-Hochberg method. Moreover, the TFBS score variability and the coefficient of variation of TF gene expression were computed. We determine the TF-archetype association when: 1) Significant variation for both TFBS score and TF gene expression and significant and positive correlation is observed (adjusted p-value <0.05 and rho>0), 2) No considerable variation is observed for TFBS score (<1Q TFBS variability) or TF gene expression (<1Q RNA coefficient of variation), independently of the correlation result, assuming that correlation will not be meaningful due to the low variability of the variables assessed. Finally, we obtained a list of archetype-TF associations that allowed us to translate archetypes to TFs and get the OCR-TF matrix (105,530 x 275) with 60,881 OCRs linked at least to one TF motif.

### Generating the Gene Regulatory Networks of B cell differentiation

Based on the different associations of OCRs with genes (cis-regulation; 62,559 links) and TFs (trans-regulation; 2,299,328 links), we were able to define the gene regulatory network (GRN) for early B cell differentiation (**Figure S2B**). To do that, from the defined OCR-gene and OCR-TF pairs, we first keep those interactions derived from the DEGs (differential expression at least in one transition), their promoter regions, and the associated enhancer elements with differential accessibility (differential chromatin accessibility at least in one transition); under the assumption that those genes and OCRs changing across cell subpopulations are going to be the ones governing the B cell differentiation process. Then, having the putative TF-OCR-gene interactions (n=1,151,024), the correlation between TF-OCR and TF-gene was computed using Spearman correlation. To reduce spurious and meaningless TF-OCR-gene interactions, we excluded from the GRN those where: 1) TF showed no differential expression (n= 469,073), 2) TF-gene association was not significantly correlated (Spearman p-value >0.05; n=186,837), 3) Meaningless interactions including (a) positive correlation for TF-gene and TF-OCR but negative correlation for TF and gene, (b) negative correlation for TF-gene and TF-OCR but positive correlation for TF and gene, (c) positive correlation for TF-gene and TF-OCR but negative correlation for OCR-gene, (d) negative correlation for TF-gene, TF-OCR and OCR-gene, and (e) positive correlation for TF-gene and OCR-gene but negative for TF-OCR (n=117,846). Finally, 377,268 TF-OCR-gene interactions were retained to reconstruct the GRN of the B cell differentiation process, involving 169 TFs, 7,310 genes, and 16,074 OCRs.

Then, the specific GRN for cell subpopulation was computed based on the chromatin accessibility and gene expression for each cell subpopulation. For example, for HSCs, the GRN captures those TF-OCR-gene interactions where genes are highly expressed in that cell subpopulation according to gene expression patterns (**Figure S6A**, genes included in brown clusters) and a median of normalized chromatin accessibility ≥ 1 (defining accessible OCR) is observed.

### Regulons cell type specificity and functionality analysis

To evaluate the cell type specificity of each predicted regulon, we calculated the Regulon Specificity Scores (RSS) per cell type and regulon. The RSS algorithm is based on the Jensen– Shannon divergence (JSD), a measure of the similarity between two probability distributions^32,33^. Specifically, we used the binary enrichment of regulon activity (based on target regions) together with the cell assignments to see how specific each predicted regulon is for each cell type. To compute the binarized regulon activity we used the GSVA score instead of the AUCell used in single-cell data. Then, a threshold of 0.2 was defined for binarizing the data, resulting in a matrix where, for each sample and regulon, we know if it is enriched (1) or not (0). Based on this binarized data and the cell type assignment for each sample we calculated the RSS using the *calculate_rss* function from scFunctions r-package [v0.0.0.9000] (https://github.com/FloWuenne/scFunctions/). The output was a data frame with RSS score for each regulon - cell type combination. The RSS was computed twice for each regulon according to TF-gene behavior, a positive correlation between TF and gene pointed to an activator role and a negative correlation to a repressor role. Heat map/dot-plot showing TF expression of the regulon on a color scale and cell-type specificity (RSS) of the regulon on a size scale (**Figure 2A, B**).

To explore the biological functions of the defined regulons we performed an over-representation analysis (ORA) of the target genes of each regulon using the *enricher* function of clusterProfiler r-package [v3.18.1]^87^. The gene ratio and −log10 from the adjusted p-value under the Benjamini-Hochberg method were computed using GO or KEGG databases (**Table S4**). Top significant and highly relevant biological pathways were selected for the summary circular plot (**Figure 2C**) generated using the *chordDiagram* function from circlize r-package [v0.4.15]^106^. For the comparison of the pathway enrichment analysis for a list of gene sets, such as the target genes of a regulon in each cell state (**Figure 3E, F** and **Figure S7C, D**), we used the *compareCluster* function from clusterProfiler r-package [v4.10.0]^107^.

### Topological properties and network distances

According to the structure of our GRNs, different network properties were computed using the igraph r-package [v1.3.5]^108^. Node betweenness (the amount of influence a node has over the flow of information in a graph through the shortest paths calculation between all pairs of nodes) and degree (the total number of edges adjacent to a node) were evaluated for the complete set of GRNs with weighted edges according to the Spearman’s coefficient of correlation between TF and gene (**Figure 2E**). Moreover, Pearson correlation was computed to compare the networks between them, and multidimensional scaling was used for visualization (**Figure 2D**).

### Single-cell samples

A total of 5 human bone marrow samples were collected for the single-cell analysis. Four samples were CD34+ human bone marrow cells, processed by single-cell multiome, and one human bone marrow sample sorted for hematopoietic lineage (sorting strategy: CD34+ CD38- CD3- CD64- CD56- CD90+; **Figure S1B**), was divided to perform both scRNA-Seq and scATAC-seq analysis.

### Single-cell Multiome Data Generation

The epigenomic landscape and gene expression of 10,000 individual nuclei from the bone marrow were examined using Single Cell Multiome ATAC + Gene Expression (10X Genomics) according to the manufacturer’s instructions (CG000365 RevC; CG000338 RevF). Briefly, 100,000 CD34+ cells were lysed for 2 min, and then 16,100 nuclei were transposed in bulk. Next, nuclei were partitioned in gel bead-in-emulsions (GEMs) in the Chromium X instrument (10X Genomics), where each nucleus was captured separately and its transposed DNA and cDNA were uniquely indexed. Pre-amplified transposed DNA/cDNA were then used to generate separate ATAC-seq and RNA-seq dual-indexed libraries. Sequencing was performed in a NextSeq2000 (Illumina) at a depth of at least 20,000 reads/nucleus for gene expression (Read1: 28; Read2: 90; i7 index: 10; i5 index: 10) and 25,000 reads/nucleus for chromatin profiling (Read1: 50; Read2: 49; i7 index: 8; i5 index: 24).

### Single-cell RNA-seq (scRNA-seq) Data Generation

The transcriptome of the hematopoietic cells from the bone marrow was examined using NEXTGEM Single Cell 3’ Reagent Kits v2 (10X Genomics), according to the manufacturer’s instructions. Between 17,600 and 20,000 cells were loaded at a concentration of 700-1,000 cells/µL on a Chromium Controller instrument (10X Genomics) to generate single-cell gel bead-in-emulsions (GEMs). In this step, each cell was encapsulated with primers containing a fixed Illumina Read 1 sequence, a cell-identifying 16 nt 10X barcode, a 10 nt Unique Molecular Identifier (UMI), and a poly-dT sequence. Upon cell lysis, reverse transcription yielded full-length, barcoded cDNA. This cDNA was then released from the GEMs, PCR-amplified, and purified with magnetic beads (SPRIselect, Beckman Coulter). Enzymatic Fragmentation and Size Selection were used to optimize cDNA size before library construction. Fragmented cDNA was then end-repaired, A-tailed, and ligated to Illumina adaptors. Final PCR amplification with barcoded primers allowed sample indexing. Library quality control and quantification were performed using Qubit 3.0 Fluorometer (Life Technologies) and Agilent’s 4200 TapeStation System (Agilent), respectively. Sequencing was performed in a NextSeq500 (Illumina) (Read1: 26 cycles; Read2:55 cycles; i7 index: 8 cycles) at an average depth of 60,226 reads/cell.

### Single-cell RNA-seq Data Analysis

The binary base call (bcl) sequence file format obtained on the 10x Genomics Chromium sequencing platform was transformed into fastq format using mkfastq from Cell Ranger 7.1.0 ^80^ (for scRNA-seq) or Cellranger-arc 2.0.2 (for multiome single-cell sequencing). The count matrix used for downstream analysis was generated using the count pipeline from Cell Ranger 7.1.0 (for scRNA-seq) or Cellranger-arc 2.0.2 (for multiome single-cell sequencing) with GRCh38 reference transcriptome provided by 10X Genomics. Following Seurat guidelines [v4.3.0.1]^109–112^, several cell and feature quality filtering were applied to each sample. First, doublets were removed using scDblFinder [v1.12.0]^113^, then, cells with >30% of mitochondrial percentage in single-cell multiome or >10% in sc RNA-seq were filtered out. Also, cells with <5% of ribosomal percentage were filtered out in the scRNA-seq sample. Finally, we filtered out extreme outlier cells or features based on lower (0.05) - upper (0.95) bound quantile for “number of features” and “counts per cell”.

Normalization of the gene expression measurements for each cell was done by multiplying the total expression by a scale factor (10,000) and log-transforming it. Feature selection criteria have been used for downstream analysis based on most variable genes, using 2,000 top genes. Feature subspaces were obtained from most variable genes using principal component analysis (PCA) using 20 components. Clustering was computed using the Louvain algorithm over principal components subspace with a resolution value of 0.5. For data visualization, UMAP was used, computing it over all PCA components obtained. Then, a first annotation based on several hematopoietic biomarkers for stem cells, lymphoid cells (T and B cells), and myeloid cells (erythroid, monocytic, and granulocytic cells) was done. Those cell clusters representing undifferentiated cells and lymphoid lineage within the CD34+ analyzed cells were kept for forward analysis. Following, to integrate the 5 single-cell transcriptional profiles, we merged the samples using Seurat. Merged samples were normalized based on the gene expression measurements for each cell by multiplying the total expression by a scale factor (10,000) and log-transforming it. Regress-out of cell cycle was applied. Feature selection criteria was used for downstream analysis based on most variable genes, using 2,000 top genes. Feature subspaces were obtained from most variable genes using principal component analysis (PCA) using 20 components. To remove the influence of dataset-of-origin we applied Harmony^114^. Clustering was computed using the Louvain algorithm over principal components subspace with a resolution value of 0.8 based on the clustree approach [v0.5.0]^115^. To annotate cell identity, we checked for cell biomarkers, cell cycle, and AUCell score [AUCell v1.20.2]^21^ derived from our bulk signatures (**Figure S8**). ^127^ To define the scRNA-seq markers for each cellular subpopulation we performed the differential expression analysis with the *FindAllMarkers* function from Seurat considering those genes present in at least 25% of cells in each identity, with an adjusted p-value < 0.01, and a logFC ≥0.25.

### Single-cell ATAC-seq Data Generation

The scATAC experiments were performed with the Chromium Next GEM Single Cell ATAC kit v1.1 from 10X Genomics following the manufacturer’s instructions. For samples with 100,000 cells or less, the nuclei isolation is done following the 10X Genomics demonstrated protocol for low cell input. Briefly, cells are collected in PBS+0.04% BSA at 4°C, and after that, cells are counted in a cell counter, and centrifugation is done to concentrate them in 50ul of PBS+0.04% BSA. After that, 45ul of supernatant is removed, and 45ul of chilled and freshly prepared lysis buffer is added. After 5 minutes of incubation on ice, 50ul of wash buffer is added. After that, centrifugation is done to remove 95ul of the supernatant without disrupting the pellet. Additional centrifugation is done with 45ul of diluted nuclei buffer to remove the supernatant. Isolated nuclei are resuspended in 7ul of chilled diluted nuclei buffer. Two of these microlitres are used to determine the cell concentration using a cell counter. The other 5ul are used immediately to generate a bulk transposition reaction following the manufacturer’s instructions and conditions. In the transposase reaction, the transposase enters the nuclei and preferentially fragments the DNA in open regions of the chromatin. In this step, adapter sequences are added to the ends of the DNA fragments. After transposition, GEMs are generated in the Chromium controller (10x Genomics). Upon GEM generation, the gel bead is dissolved. Oligonucleotides containing an Illumina P5 sequence, a 16nt 10X barcode, and a read1 sequence are released and mixed with DNA fragments and master mix. After incubation, GEMs are broken, and pooled fractions are recovered. P7, a sample index, and read2 sequence are added during library construction via PCR. Library quality control and quantification were performed using Qubit 3.0 Fluorometer (Life Technologies) and Agilent’s 4200 TapeStation System. Bioanalyzer 2100 (Agilent). Sequencing was performed using a NextSeq500 (Illumina) (Rd1 50 cycles, i7 Index 8 cycles, i5 index 16 cycles, and Rd2 50 cycles) to an average depth of 14,900 fragments/cell.

### Single-cell ATAC-seq Data Analysis

The bcl file format obtained on the 10x Genomics Chromium sequencing platform was, in this case, transformed into fastq format using mkfastq from Cell Ranger ATAC 1.1.0 ^116^ (for scRNA-seq) or Cellranger-arc 2.0.2 (for multiome single-cell sequencing). The count matrix used for downstream analysis was generated similarly using the count pipeline from Cell Ranger ATAC 1.1.0 (for scATAC-seq) or Cellranger-arc 2.0.2 (for multiome single-cell sequencing) with GRCh38 reference transcriptome provided by 10X Genomics.

To make bulk and single-cell ATAC-seq comparable, a re-quantified version of the scATAC-seq original count matrix was conducted to generate a new feature x cell raw count matrix with the same OCRs from bulk as feature space using the *FeatureMatrix* function from Signac package [v1.9.0]^117^. The new scATAC-seq matrix contains recalculated count values based on scATAC-seq genomic fragments according to bulk ATAC-seq feature space. Then, we implemented quality control criteria, retaining cells with a Fraction of Reads in Peaks (FRIP) value of at least 50% and a Transcription Start Site (TSS) enrichment score above 5. Cells with a log-transformed number of unique fragments per barcode below 3.8 were filtered out. Furthermore, doublet cells were identified and excluded using the Scrublet algorithm with default parameters ^118^. Dimensionality reduction was conducted using Latent Dirichlet Allocation (LDA) by performing Gibbs sampling for various topic counts (K). The optimal K value was selected based on the highest consensus scores from four pycisTopic proposed metrics^119^.

Accessibility imputation was carried out using the *impute_accessibility* function from pycisTopic package [v1.0.3.]^20^, with a scaling factor of 10^6^ to normalize accessibility scores across selected cells and regions. Subsequently, gene activity scores were computed using the *get_gene_activity* function from pycisTopic package, incorporating region-cell probabilities, distance weights, and upstream/downstream search spaces ranging from 1 kb to 100 kb to refine gene activity assignments. Distance weights and other parameters, such as gene boundaries and TSS extensions, default values were employed to optimize the accuracy of the activity predictions.

Label transfer was executed in two phases. First, annotations derived from scRNA-seq were transferred to the multiome scATAC-seq (activity scores) samples, employing annotation transfer methods including Harmony, BBKNN, Scanorama, and Canonical Correlation Analysis (CCA). The accuracy of these transfers was evaluated against the annotations of the corresponding cell labels in the scRNA-seq, with Harmony yielding the highest accuracy across four multiome samples. In the second phase, this transfer method was utilized to annotate the multiome samples and the unpaired scATAC-seq sample.

Finally, samples were integrated using the Harmony method^114^, facilitating the generation of a common subspace for visualization.

### Single-cell ATAC-seq Regulon Analysis

To explore enhancer gene regulatory networks (eGRNs), we integrated the previously identified scATAC-seq data with guidelines suggested by the SCENIC+ workflow [v1.0.1]^20^. In line with SCENIC+ recommendations, we identified candidate enhancer regions by binarizing region-topic probabilities and analyzing differentially accessible regions (DARs). The binarization process used Otsu’s method, followed by dropout imputation by multiplying cell-topic and topic-region distributions, as previously established by cisTopic^119^. Normalization of the data was conducted using a scale factor of 10^4^, after which highly variable peaks were identified. DARs were computed using the Wilcoxon rank-sum test, maintaining the same ordered labeling of cell types that was established in the prior scRNA-seq analysis.

To generate eGRNs, we merged scRNA-seq and scATAC-seq data to create a pseudo-multiome dataset. For each cell type annotation, we randomly selected cells from both datasets. Meta-cells were constructed by averaging scRNA-seq and scATAC-seq counts for each annotation with K = 5 cells. Motif enrichment analysis was then performed using the SCENIC+ pipeline, integrating binarized topics and DARs from pycisTopic [v1.0.3]. We applied the cisTarget and Differential Enrichment of Motifs (DEM) algorithms for motif discovery.

CisTarget identified genomic regions from the SCREEN database (HG38) using the Cluster-Buster tool [v3.1.0]^120^ and ranked motifs by motif scores. A minimum overlap of 40% between input regions and reference motifs was required. Recovery curve-based scoring and Normalized Enrichment Scores (NES) were computed for each motif, retaining only motifs with an NES > 3. DEM performed a Wilcoxon test to assess motif enrichment between foreground and background regions. Motifs with adjusted p-values < 0.05 (Bonferroni corrected) and a log2FC > 0.5 were selected.

Cistromes were generated by merging motif-based regions associated with transcription factors (TFs), providing a comprehensive map of TF-target regulatory interactions. Finally, region-to-gene distances were computed, and TF-target importances were evaluated using the Gradient Boosting Machine (GBM) method, adhering to SCENIC+ guidelines.

### Public RNA-Seq data from BCP ALL patients

Transcriptomic data from BCP ALL patients (leukemic cells from bone marrow aspirates) were obtained from the European Genome-phenome Archive. The RNA-Seq data (BAM files) from EGAD00001002151, EGAD00001002692, and EGAD00001002704 datasets were downloaded after the corresponding access request. Only data from pediatric individuals belonging to a well-known molecular subgroup^121^, ranging from G1 to G8 (MEF2D fusions, TCF3–PBX1, ETV6– RUNX1/-like, DUX4 fusions, ZNF384 fusions, BCR–ABL1/Ph-like, Hyperdiploidy, KMT2A fusions) were included in the analysis, resulting in 356 patients (**Table S12**).

As BAM files were mapped to GRCh37, we first obtained the Fastq files using the *bamtofastq* function from bedtools [version 2.25.0] ^95^. Next, the read quality was assessed with FastQC software [version 0.11.7] (Babraham Institute). Low quality, short reads, and TruSeq3-SE adapters were trimmed using Trimmomatic [version 2.6.0a] ^82^ under the following parameters: ILLUMINACLIP:TruSeq3-SE:2:30:10 LEADING:3 TRAILING:3 SLIDINGWINDOW:4:15 MINLEN:75. Reads were then mapped to the December 2013 Homo sapiens high coverage assembly GRCh38 from the Genome Reference Consortium GRCh38 (hg38) using STAR [version 2.6.0a]^83^ and counted for each gene using htseq-count^84^ with –m intersection-nonempty option and a GTF file from GRCh38 Ensembl annotation release 96, getting the information for a total of 58,884 genes. Genes with a minimum read count of 1 in all population replicates and an associated ensemble ID were retained (25,914 genes). Next, the batch correction was done using the *ComBat_seq* function from sva r-package [v3.38.0] ^122,123^. Counts were normalized under TMM normalization and voom transformation from edgeR r-package [v3.32.1]^85^. Multidimensional scaling (MDS) was used for data exploration. Differential expression analysis (DEA) of leukemic subgroups was performed using limma. Those genes with low coefficient of variation (<25% cv) were filtered out before DEA to win statistical power. Genes with an adjusted p-value ≤ 0.01 (Benjamini-Hochberg) and absolute fold-change ≥ 1.5 at least in 4 contrasts were defined as significant (subgroup biomarkers). The subgroup biomarkers list can be found in **Table S12**.

### Over-representation analysis in public BCP ALL data

The over-representation analysis (ORA) of the gene sets from B-ALL subgroups (**Table S12**) was performed in the biomarkers of the different B-cell populations (for both, bulk and single-cell approaches) using the *enricher* function of clusterProfiler r-package [v3.18.1]^87^ (**Figure 5B, F**). The gene ratio and −log10 from the adjusted p-value under the Benjamini-Hochberg method were computed. The ORA was also applied to explore the enrichment of 711 binding sites defined as the highest affinity binding sites for ETV6-RUNX1 fusion under ChiP-Seq analysis^55^ in the consensus OCRs for each cell subpopulation (**Figure 5D**). Finally, we applied ORA to explore if there was a significant enrichment of the regulons identified along B-cell differentiation in the different B-ALL subgroups (**Figure 5E, G**).

### Public ChIP-Seq Histone marks

ChIP-Seq data from ENCODE project (https://www.encodeproject.org/)^124^ for H3K4me1, H3K4me3, and H3K27ac on B cells, naive B cells, and common myeloid progenitors CD34+ was downloaded (**Table S1**). To get the consensus regions for each cell subpopulation and histone mark, we found the genome regions that are covered by at least n-1 of the sets of peaks (n being the total number of samples for a specific cell subpopulation), and then we merged the nearby peaks with a min gap width of 31. To get the consensus regions for a histone mark in the whole population, we get the regions in the genome where coverage is at least in 1 of the consensus histone marks from the different cell subpopulations and then merged the nearby peaks with a min gap width of 31, resulting in 166,295 H3K4me1 regions, 37,952 H3K4me3 regions, and 159,079 H3K27ac regions. These processes were done using the rtracklayer [v1.58.0] and GenomicRanges [v1.50.2] r-packages ^92,93^. Then, the overlap between our OCRs and histone marks was assessed using the *findOverlaps* function (minoverlap=6) of GenomicRanges (**Figure S11A**).

### Public Hi-C data

Hi-C public data of naive B cells were obtained from EGAD00001006486^60^.

The valid reads of Hi-C experiments were processed with TADbit version 1.1 ^125^. Hi-C data were normalized using the *oned* function of dryhic r-package [v0.0.0.9100] ^126^ at 100 kb of resolution to remove known experimental biases, a method that controls for the presence of abnormal karyotypes in cancer samples. Topologically associating domains (TADs) were detected at 40 kbp resolution using the TopDom algorithm ^127^. The significant Hi-C interactions were called with the Fit-Hi-C tool [v1.1.3] ^128,129^, binned at 10 kb of resolution and with the default thresholds; p-value< 0.001 and q-value<0.001.

To study the compartmentalization of the genome in naive B cells, we used merged biological replicates resulting in interaction Hi-C maps with around 300 million valid reads each. Briefly, normalized chromosome-wide interaction matrices at 100 kb resolution were transformed into Pearson correlation matrices. These correlation matrices were then used for PCA, for which the first eigenvector (EV) normally delineates genome segregation. The multi-modal distribution of the EV coefficients from the B cells dataset was modeled as a Gaussian mixture with three components (k = 3). To estimate the mixture distribution parameters, an Expectation-Maximization algorithm using the *normalmixEM* function from the mixtools r-package [v1.2.0] was applied ^130^. A Bayesian Information Criterion (BIC) was computed for the specified mixture models of clusters (from 1 to 10) using the *mclustBIC* function from mclust r-package [v5.4.9] ^86^. Three underlying structures were defined; alternative compartmentalization into A-type (with the most positive EV values), B-type (with the most negative EV values), and I-type (an intermediate-valued region with a different distribution) compartments. Two intersection values were defined at the intersection points between two compartments and used as standard thresholds to categorize the data into the three different compartments; that is: the A-type compartment was defined for EV values >= +0.1388, the I-type compartment was defined for EV values between +0.1388 and −0.01955, and the B-type compartment was defined for EV values between <= −0.01955.

Then, we checked the overlap between our OCRs and the compartments identified using the *findOverlaps* function (default parameters) of GenomicRanges r-package [v1.50.2] (**Figure S11B**). When an OCR overlaps with two compartments, we assigned it to the compartment with a higher overlap.

### Public promoter capture Hi-C interaction data

Detected interactions using promoter capture Hi-C (PCHiC) technology from Javierre et al.^61^ were downloaded. From the PCHiC peak matrix, we select all those interactions with a CHiCAGO score ≥ 5 in the naive B cell population, resulting in 197,442 interactions. The coordinates of the interacting regions were translated from GRCh37 to GRCh38 using liftOver r-package [v1.18.0]. Then, we check the overlap between our interactions (OCR-gene associations) and their interactions using the *findOverlaps* function (minoverlap=6) of GenomicRanges r-package [v1.50.2] (**Figure S11C** showing some of the overlapping interactions).

### Web application

Shiny r-package [v1.8.0]^131^ was used to generate a user-friendly web interface for browsing called B-rex (https://translationalbio.shinyapps.io/brex/). For single-cell RNA-seq the ShinyCell template [v2.1.0] was used^132^. The Shiny app is hosted on the https://www.shinyapps.io/.

## DATA AND SOFTWARE AVAILABILITY

The EGA accession number for gene expression and chromatin accessibility data reported in this paper is EGAS00001007296.

## ACKNOWLEDGEMENTS

This work was supported by the Instituto de Salud Carlos III and co-financed by ERDF A way of making Europe (PI17/00701, and PI20/01308), CIBERONC (CB16/12/00489) and RICORS TERAV (RD21/0017/0009); Gobierno de Navarra (AGATA 0011-1411-2020-000010/0011-1411-2020-000011; Fundación La Caixa (GR-NET NORMAL-HIT HR20-00871); and Cancer Research UK [C355/A26819], FC AECC and AIRC under the Accelerator Award Program. The Multiple Myeloma Research Foundation Networks of Excellence 2017 Immunotherapy Program Grant Award, the International Myeloma Foundation (Brian van Novis), the Paula and Rodger Riney Foundation.

N.P.P was funded by a Juan de la Cierva-formación fellow (FJC2019-042304-I) from the Spanish Ministry of Science and Innovation (MCIN) and by a Ramón y Cajal fellow (RYC2021-032197-I) from the MCIN/AEI/10.13039/501100011033 and European Union “NextGenerationEU”/PRTR. The contribution from DGC and JT was partially funded from King Abdullah University of Science and Technology

This study makes use of data generated by the St. Jude Children’s Research Hospital – Washington University Pediatric Cancer Genome Project (EGAD00001002692 and EGAD00001002704), and data generated by IDIBAPS (EGAD00001006486).

We particularly acknowledge the healthy donors for their participation in this study, and the Biobank of the University of Navarra for its collaboration.

## Supplementary Figure legends

**Figure S1. Gating strategy of B-cell subpopulations. (A)** Gating strategy followed in the flow cytometry analysis to obtain the different B cell subpopulations from bone marrow samples, shown for 1 representative sample. **(B)** Gating strategy followed in the flow cytometry analysis to isolate the CD34+ single-cell samples from the bone marrow, shown for 1 representative sample.

**Figure S2. Data generated and computation analysis followed to define the GRN of B-cell differentiation derived from chromatin accessibility and gene expression coordination. (A)** Schematic representation of data generated. Bone marrow and peripheral blood samples were collected from 13 healthy individuals and 8 B-cell subpopulations isolated by flow cytometry. Then, each sample underwent RNA-seq and ATAC-seq profiling, recovering 79 and 78 high-quality samples, respectively. Individual omics analyses were performed to assess the differential gene expression in RNA-seq data and the genomic characterization, differential accessibility, and TF binding site activity in ATAC-seq. **(B)** Illustration of the computational workflow followed to decipher the GRN from the RNA-seq and the ATAC-seq coordination. We first defined the GRN according to three main steps**: I) *Cis-regulatory elements (CREs)***. To determine the cis-regulatory elements (CREs) of a gene, we follow two approaches: 1) For promoter OCRs, we directly annotate OCRs to genes based on TSS distance (<1kb), and 2) For intronic or distal intergenic OCRs, we explored the association between chromatin accessibility and gene expression within a window of ±1Mb from the gene’s TSS. **II) *Trans-regulatory elements (TFs)***. To annotate OCRs to putative TF binding motifs, we first identify the footprints within our OCRs. Next, we annotated the putative transcription factor binding motif to each footprint using the Vierstra database, resulting in an OCR-TF matrix (105,530 x 275) with 60,881 OCRs linked at least to one TF motif. ***III) TF-OCR-gene interactions (GRNs)***. Based on the different associations of OCRs with genes (cis-regulation; 62,559 links) and TFs (trans-regulation; 2,299,328 links), we were able to define the gene regulatory network (GRN) for early B cell differentiation showing 1,823,177 TF-OCR-gene putative interactions. Next, we curated the GRN to highlight the differentiation program. To do that, we first keep those interactions derived from the DEGs, their promoter regions, and the associated enhancer elements with differential accessibility. Then, we reduced spurious and meaningless TF-OCR-gene interactions based on TF-OCR and TF-gene coordination. Finally, 377,268 TF-OCR-gene interactions were retained to reconstruct the GRN of the B cell differentiation process, involving 169 TFs, 7,310 genes, and 16,074 OCRs. Finally, the specific GRN for each cell subpopulation was computed based on the chromatin accessibility and gene expression. All these results have been provided as a Shiny app, called B-rex, publicly available at: https://translationalbio.shinyapps.io/brex/.

**Figure S3. TF activity profile across B-cell differentiation.** Heatmap representation of the TFBS activity score profiling for all the TF identified (n=275). Samples are shown by columns (n=78) and grouped by cell subpopulation, and TFs by rows.

**Figure S4. Transcriptional signature of human B-cell subpopulations. (A)** Heatmap representation of 8,642 genes defining the signature profile of the B-cell subpopulations. Biomarker genes for each B-cell subpopulation were derived from significant genes (logFC≥1.5 and adjusted p-value≤0.05) in at least 4 contrasts (see methods). The Z-score of normalized gene expression levels is represented. Up-regulated genes are shown in red and down-regulated genes in blue. Each row shows an individual gene and each column an individual sample. Color coding according to cell subpopulation is shown on the top row of the heatmap. **(B)** Upset plot showing the overlap between the gene expression profile of each B-cell subpopulation.

**Figure S5. Transcriptional profile of public mice B-cell subpopulations (A)** Principal Component Analysis (PCA) of gene expression from public mice B-cell subpopulations (17 samples) showing the distribution of cell subpopulations. **(B)** Heatmap representation of gene expression profile for 54 well-known genes associated with mice B-cell lymphopoiesis. Samples are shown by columns (n=17; grouped by cell subpopulation), and genes by rows (n=58; grouped by biological processes). Normalized gene expression levels are represented.

**Figure S6. Profiling of those genes, chromatin regions, and TF constructing the Gene Regulatory Network of early B cell differentiation. (A)** Heatmap representation of genes included in the GRN (n=7,310 genes). On the left is the Z-score of gene expression values. On the right is a binary representation of gene expression patterns, with high expression in brown. Samples are shown by columns and grouped by cell subpopulation. **(B)** Heatmap representation of OCRs included in the GRN (n=16,074 OCRs). On the left is the Z-score of chromatin accessibility values. In the middle is the genomic annotation of each OCR (promotor, intronic, or distal intergenic). On the right is a binary representation of the chromatin accessibility patterns, with high accessibility in brown, and regions with no significant changes in chromatin accessibility in gray, all corresponding to promoter regions. Samples are shown by columns and grouped by cell subpopulation. **(C)** Heatmap representation of the TFBS activity score of those TFs included the GRN (n=169 TFs). Samples are shown by columns and grouped by cell subpopulation. **(D)** Heatmap showing the TFs that are included in each cell-specific GRN. TFs are shown by columns (n=169) and the B-cell-specific GRNs by row (n=8).

**Figure S7. EBF1 regulon. (A)** Heatmap representation of gene expression for TF regulators of EBF1. Samples are shown by columns (grouped by cell subpopulation) and TFs by rows. Normalized gene expression levels are represented. On the right side rho values of Spearman correlation for each TF vs EBF1 are shown. On the top, a barplot representation of the EBF1 gene expression is shown. **(B)** Heatmap representation of gene expression for TF regulated by EBF1. Samples are shown by columns (grouped by cell subpopulation) and TFs by rows. Normalized gene expression levels are represented. On the right side rho values of Spearman correlation for each TF vs EBF1 are shown. On the top, a barplot representation of the EBF1 gene expression is shown. **(C)** Representation of the enriched GO terms derived from the pathway analysis of the EBF1-regulon-activated genes and **(D)** EBF1-regulon-repressed genes for each csGRN. The adjusted p-value is shown in a red-blue color range for each term and the number of genes enriched in each term (counts) as dot size.

**Figure S8. Single-cell RNA-seq annotation and regulon identification. (A)** Single-cell RNA-seq Uniform Manifold Approximation and Projection (UMAP) dimensionality reduction of 28,891 human CD34+ cells drawing the B-cell lymphopoiesis based on bulk HSC, CLP, and pro-B gene signatures. The AUCell score for each signature is represented. **(B)** Dot plot showing the gene expression of the B-cell markers used for annotating the individual single-cell RNA-seq clusters. Average gene expression is shown in a color range and the percentage of expressing cells within a cluster by dot size. **(C)** Single-cell RNA-seq Uniform Manifold Approximation and Projection (UMAP) dimensionality reduction of 28,891 human CD34+ cells drawing the B-cell lymphopoiesis based on the cell cycle phase of each cell (red: G1; blue: S phase; and green: G2/M phase). **(D)** Scatter plot showing the number of target regions versus TF expression-to-region AUC Pearson correlation coefficients for each eRegulon inferred by SCENIC+. eRegulons are selected based on a threshold on the correlation coefficient, indicated by the dotted lines (−0.75 and 0.7).

**Figure S9. Transcriptional signature of human B-ALL subtypes and B-ALL origin inference based on csGRNs. (A)** Heatmap representation of 5,749 genes defining the signature profile of the B-ALL subtypes. Biomarker genes for each B-ALL subtype were derived from significant genes (logFC≥1.5 and adjusted p-value≤0.05) in at least 4 contrasts (see methods). The Z-score of normalized gene expression levels is represented. Up-regulated genes are shown in red and down-regulated genes in blue. Each row shows an individual gene and each column an individual sample. Color coding according to cell subpopulation is shown on the top row of the heatmap. **(B)** Upset plot showing the overlap between the gene expression profile of each B-ALL subtype. **(C)** Over Representation Analysis (ORA) of B-ALL subtypes signatures over the B-cell csGRNs signatures. The dot size represents the −log10(adjusted p-value) and the color range the gene ratio.

**Figure S10. Sample variability explained by transcriptional and chromatin accessibility profiling. (A)** Heatmap representation of Pearson correlation matrices (rho) between samples based on RNA-seq data (top left), ATAC-seq data (top right), ATAC-seq data at all promoter OCRs (bottom left), ATAC-seq data at all intronic OCRs (bottom center), and ATAC-seq data at all distal intergenic OCRs (bottom right). The corresponding OCR proportion of each OCR-type (promoter, intronic, distal intergenic) is shown. Color-coding according cell subpopulation is shown on the top row of each heatmap. **(B)** A t-SNE representation of OCRs identified in the study. First, Gini index for all OCRs; Gini index highlights OCR variability. OCRs with large variability are shown in blue and those with low variability in red. Following OCR distribution for promoter OCRs, intronic OCRs and distal intergenic OCRs. High density regions are shown in yellow and low density in dark blue. **(C)** Variance component decomposition of the mRNA expression for every gene (as column), in a variance component model that discretizes the explanatory power of promoter OCRs (in green), distal enhancer OCRs (in blue), and unexplained variance (in red). The proportion of variance explained has been shown for the total number of identified genes (top left), for differentially expressed genes (bottom left), and for the genes not differentially expressed (bottom right). Also, the variance components model results for two permuted matrices per gene model are shown (top right).

**Figure S11. Enrichment of histone marks and A-type compartments of chromatin conformation into the identified CREs. (A)** Barplot representation of OCRs overlapping histone marks; shown as proportions. Distribution for all OCRs, OCRs defined as candidate cis-regulatory elements (cCREs), and OCRs not defined as candidate cis-regulatory elements (cCREs) are shown. The proportion of OCRs without histone marks information are shown in pink (no histone mark). **(B)** Barplot representation of OCRs overlapping Hi-C compartments; shown as proportions. Distribution for all OCRs, OCRs defined as candidate cis-regulatory elements (cCREs), and OCRs not defined as candidate cis-regulatory elements (cCREs) are shown. Hi-C compartments are differentiated between active compartment (A, in coral), intermediate compartment (I, in blue), and inactive compartment (B, in green). The proportion of OCRs without compartment information are shown in gray (missing). **(C)** Heatmap representation of 2,161 OCR-gene pairs overlapping with Promoter Capture-HiC public interactions. The left heatmap shows the chromatin accessibility levels (in z-score) and the right heatmap the gene expression level (in z-score). Each row corresponds to an OCR-gene pair. Color coding according to cell subpopulation is shown on the top row of the heatmap. Some OCR-gene pairs are highlighted on the right.

