## Supplementary Figures for "Uncovering the regulatory landscape of early human B-cell lymphopoiesis and its implications in the pathogenesis of B-cell acute lymphoblastic leukemia"

### Figure S1

A

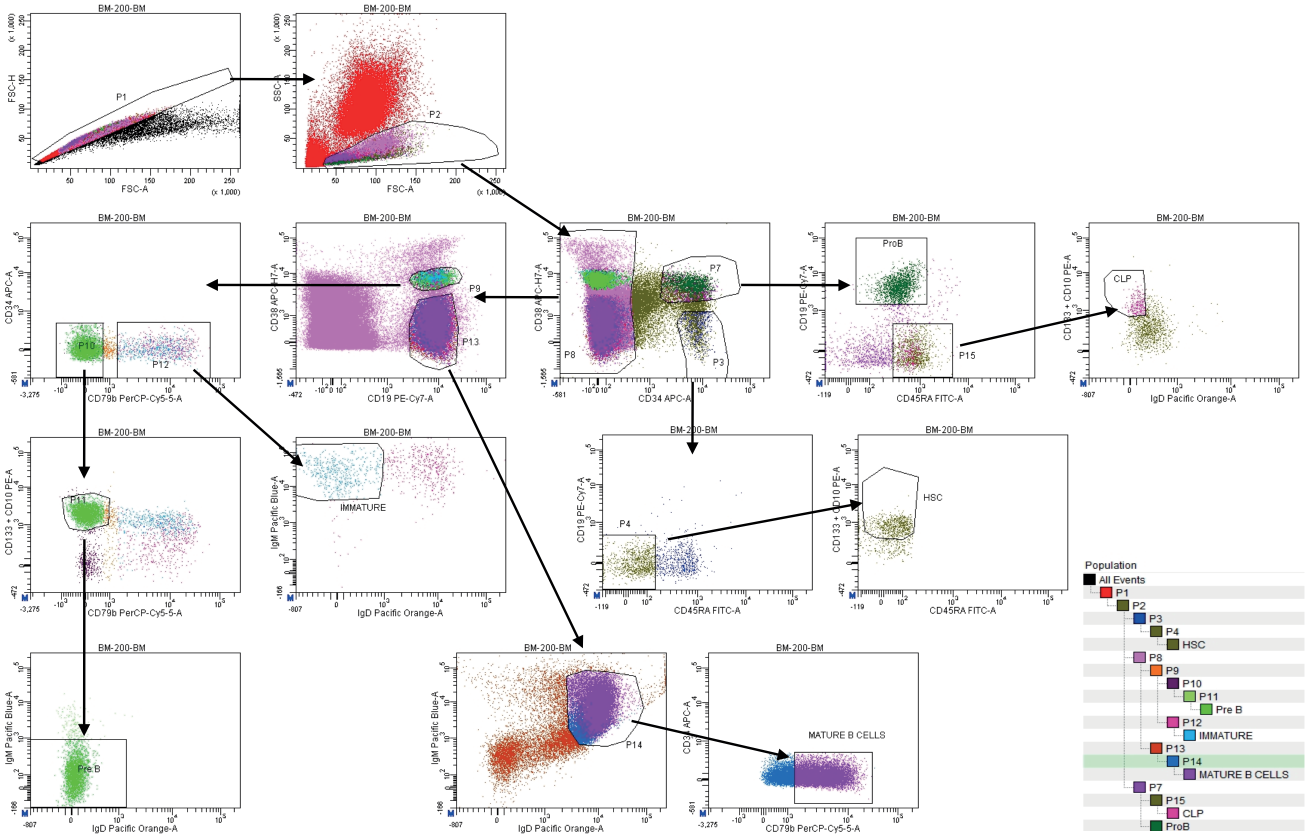
BM-200-BM
P1
P2
P3
P4
P7
P8
P9
P10
P11
P12
P13
P14
P15
ProB
CLP
HSC
IMMATURE
Pre B
MATURE B CELLS
FSC-H
FSC-A
SSC-A
CD34 APC-A
CD38 APC-H7-A
CD19 PE-Cy7-A
CD79b PerCP-Cy5-5-A
CD45RA FITC-A
CD133 + CD10 PE-A
IgD Pacific Orange-A
IgM Pacific Blue-A
Population
All Events
P1
P2
P3
P4
HSC
P8
P9
P10
P11
Pre B
P12
IMMATURE
P13
P14
MATURE B CELLS
P7
P15
CLP
ProB

B

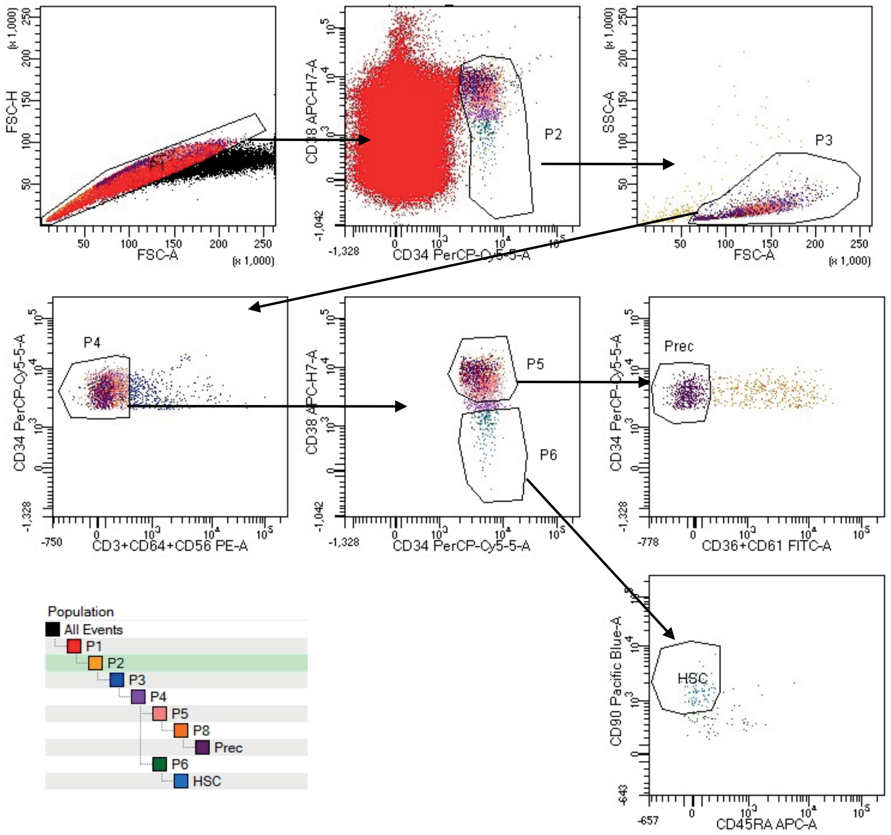
P1
P2
P3
P4
P5
P6
Prec
HSC
FSC-H
FSC-A
SSC-A
CD38 APC-H7-A
CD34 PerCP-Cy5-5-A
CD3+CD64+CD56 PE-A
CD36+CD61 FITC-A
CD90 Pacific Blue-A
CD45RA APC-A
Population
All Events
P1
P2
P3
P4
P5
P8
Prec
P6
HSC

Figure S2

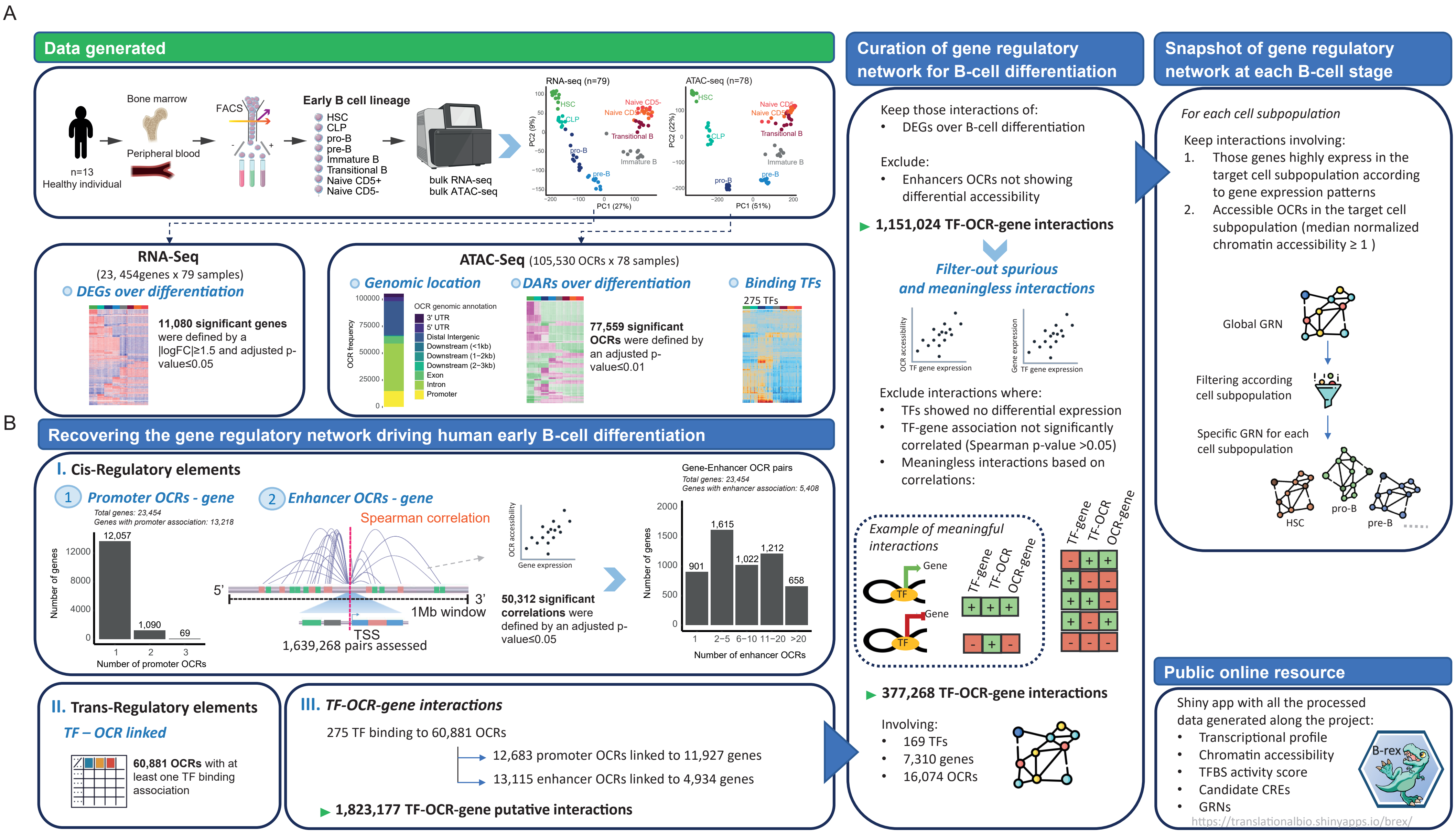
A
Data generated
Bone marrow
Peripheral blood
n=13
Healthy individual
FACS
Early B cell lineage
HSC
CLP
pro-B
pre-B
Immature B
Transitional B
Naive CD5+
Naive CD5-
bulk RNA-seq
bulk ATAC-seq
RNA-seq (n=79)
PC1 (27%)
PC2 (9%)
ATAC-seq (n=78)
PC1 (51%)
PC2 (22%)
RNA-Seq
(23, 454genes x 79 samples)
DEGs over differentiation
11,080 significant genes were defined by a |logFC|≥1.5 and adjusted p-value≤0.05
ATAC-Seq (105,530 OCRs x 78 samples)
Genomic location
DARs over differentiation
Binding TFs
OCR genomic annotation
3' UTR
5' UTR
Distal Intergenic
Downstream (<1kb)
Downstream (1–2kb)
Downstream (2–3kb)
Exon
Intron
Promoter
OCR frequency
77,559 significant OCRs were defined by an adjusted p-value≤0.01
275 TFs
Curation of gene regulatory network for B-cell differentiation
Keep those interactions of:
• DEGs over B-cell differentiation
Exclude:
• Enhancers OCRs not showing differential accessibility
1,151,024 TF-OCR-gene interactions
Filter-out spurious and meaningless interactions
OCR accessibility
TF gene expression
Gene expression
Exclude interactions where:
• TFs showed no differential expression
• TF-gene association not significantly correlated (Spearman p-value >0.05)
• Meaningless interactions based on correlations:
Example of meaningful interactions
Gene
TF
TF-gene
TF-OCR
OCR-gene
377,268 TF-OCR-gene interactions
Involving:
• 169 TFs
• 7,310 genes
• 16,074 OCRs
Snapshot of gene regulatory network at each B-cell stage
For each cell subpopulation
Keep interactions involving:
1. Those genes highly express in the target cell subpopulation according to gene expression patterns
2. Accessible OCRs in the target cell subpopulation (median normalized chromatin accessibility ≥ 1 )
Global GRN
Filtering according cell subpopulation
Specific GRN for each cell subpopulation
HSC
pro-B
pre-B
B
Recovering the gene regulatory network driving human early B-cell differentiation
I. Cis-Regulatory elements
1 Promoter OCRs - gene
Total genes: 23,454
Genes with promoter association: 13,218
12,057
1,090
69
Number of genes
Number of promoter OCRs
2 Enhancer OCRs - gene
Spearman correlation
5'
3'
1Mb window
TSS
1,639,268 pairs assessed
OCR accessibility
Gene expression
50,312 significant correlations were defined by an adjusted p-value≤0.05
Gene-Enhancer OCR pairs
Total genes: 23,454
Genes with enhancer association: 5,408
901
1,615
1,022
1,212
658
Number of genes
Number of enhancer OCRs
II. Trans-Regulatory elements
TF – OCR linked
60,881 OCRs with at least one TF binding association
III. TF-OCR-gene interactions
275 TF binding to 60,881 OCRs
12,683 promoter OCRs linked to 11,927 genes
13,115 enhancer OCRs linked to 4,934 genes
1,823,177 TF-OCR-gene putative interactions
Public online resource
Shiny app with all the processed data generated along the project:
• Transcriptional profile
• Chromatin accessibility
• TFBS activity score
• Candidate CREs
• GRNs
B-rex
https://translationalbio.shinyapps.io/brex/

Figure S3

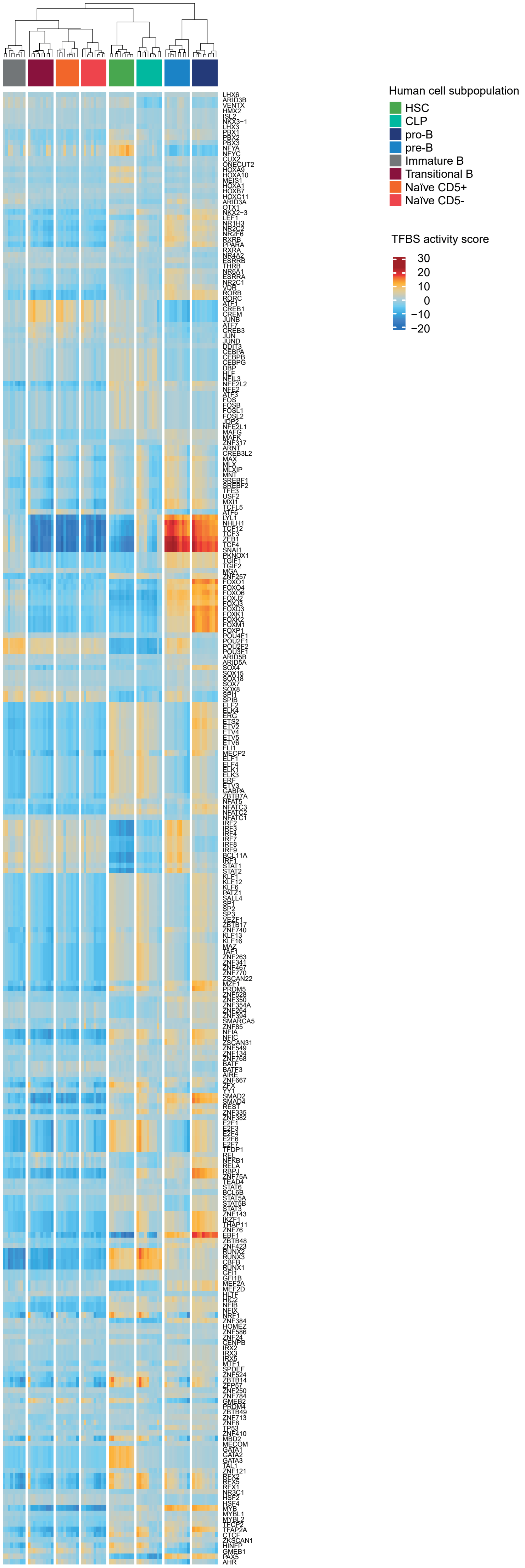
Human cell subpopulation
HSC
CLP
pro-B
pre-B
Immature B
Transitional B
Naïve CD5+
Naïve CD5-
TFBS activity score
30
20
10
0
−10
−20

### Figure S4

A

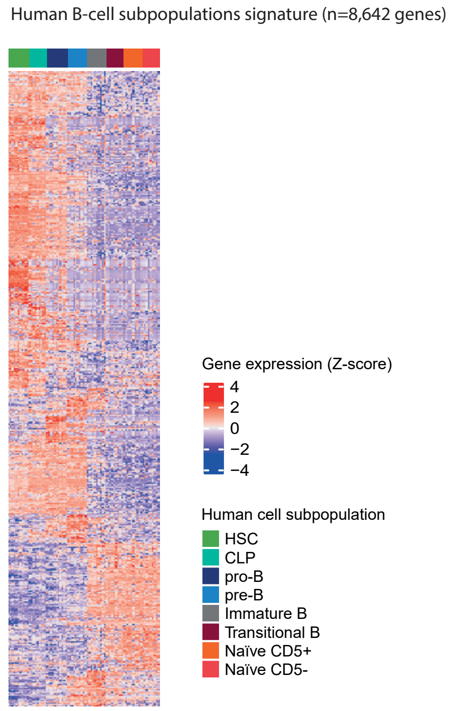
Human B-cell subpopulations signature (n=8,642 genes)
Gene expression (Z-score)
4
2
0
−2
−4
Human cell subpopulation
HSC
CLP
pro-B
pre-B
Immature B
Transitional B
Naïve CD5+
Naïve CD5-

B

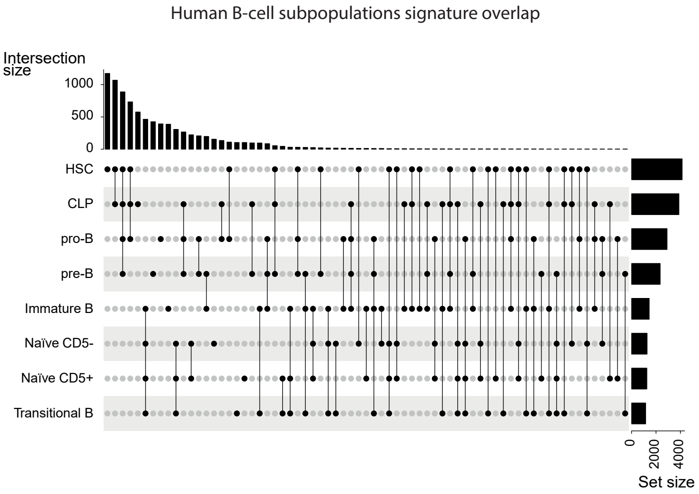
Human B-cell subpopulations signature overlap
Intersection size
1000
500
0
HSC
CLP
pro-B
pre-B
Immature B
Naïve CD5-
Naïve CD5+
Transitional B
0
2000
4000
Set size

Figure S5

A

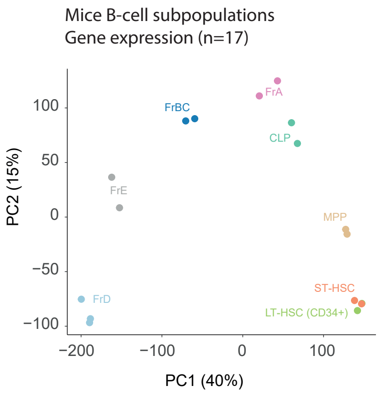
Mice B-cell subpopulations
Gene expression (n=17)
PC2 (15%)
PC1 (40%)
100
50
0
−50
−100
−200
−100
0
100
FrBC
FrA
CLP
FrE
MPP
FrD
ST-HSC
LT-HSC (CD34+)

B

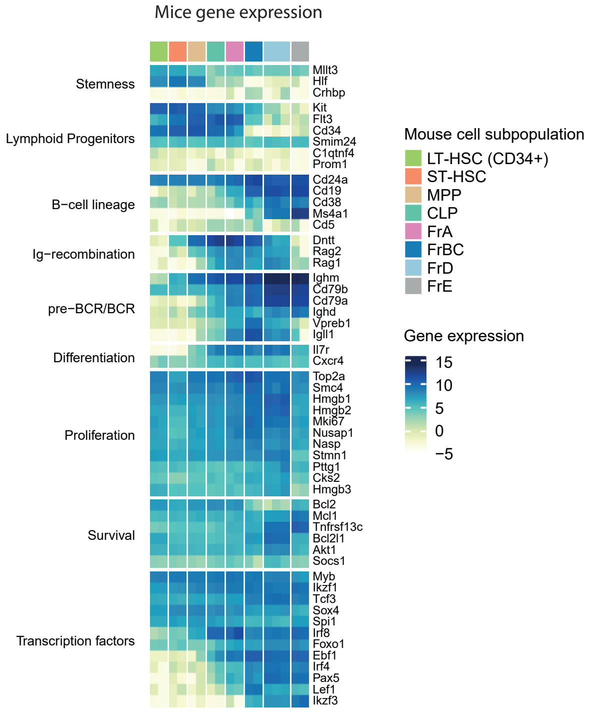
Mice gene expression
Stemness
Mllt3
Hlf
Crhbp
Lymphoid Progenitors
Kit
Flt3
Cd34
Smim24
C1qtnf4
Prom1
B−cell lineage
Cd24a
Cd19
Cd38
Ms4a1
Cd5
Ig−recombination
Dntt
Rag2
Rag1
pre−BCR/BCR
Ighm
Cd79b
Cd79a
Ighd
Vpreb1
Igll1
Differentiation
Il7r
Cxcr4
Proliferation
Top2a
Smc4
Hmgb1
Hmgb2
Mki67
Nusap1
Nasp
Stmn1
Pttg1
Cks2
Hmgb3
Survival
Bcl2
Mcl1
Tnfrsf13c
Bcl2l1
Akt1
Socs1
Transcription factors
Myb
Ikzf1
Tcf3
Sox4
Spi1
Irf8
Foxo1
Ebf1
Irf4
Pax5
Lef1
Ikzf3
Mouse cell subpopulation
LT-HSC (CD34+)
ST-HSC
MPP
CLP
FrA
FrBC
FrD
FrE
Gene expression
15
10
5
0
−5

Figure S6

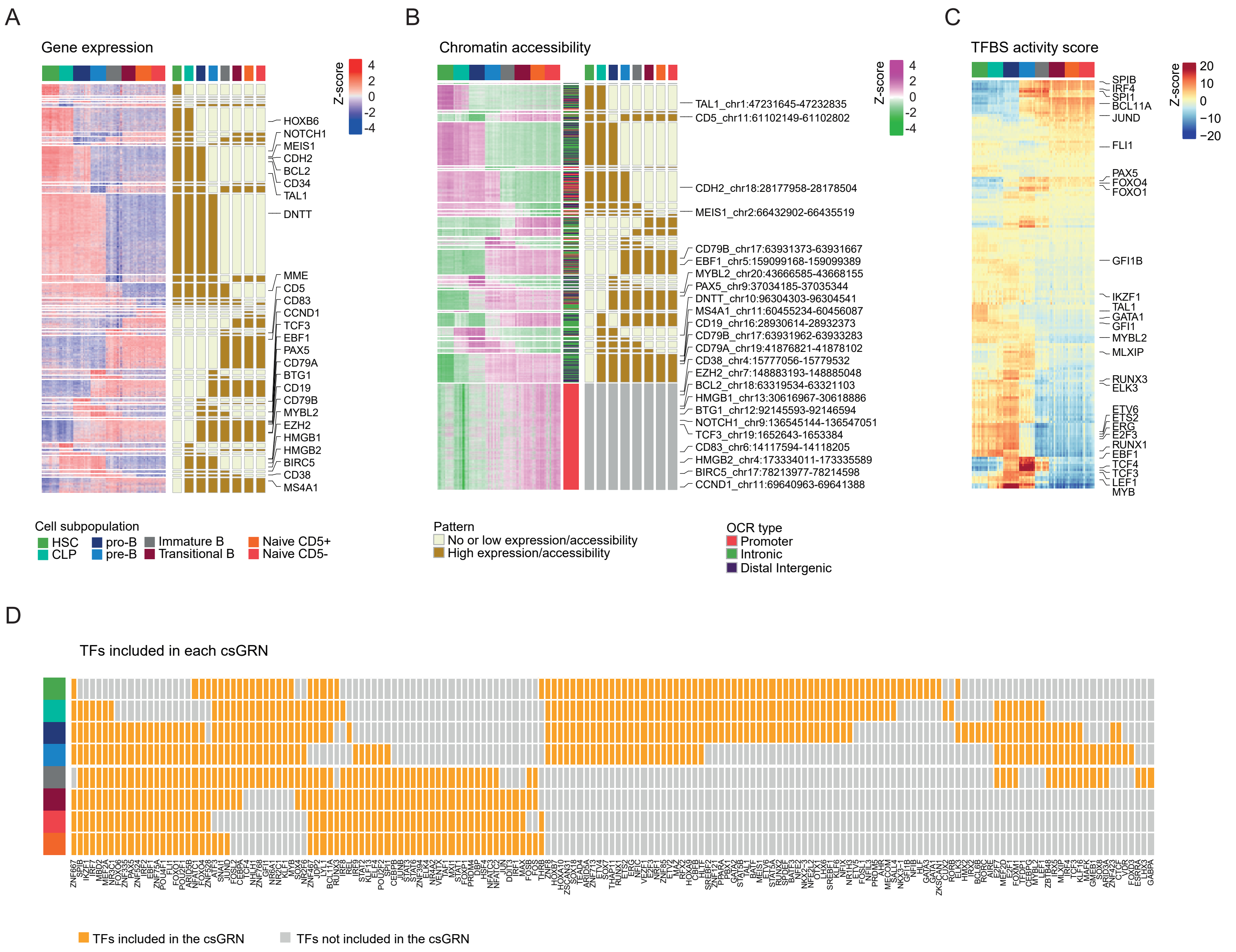
A
Gene expression
Z-score
4
2
0
-2
-4
HOXB6
NOTCH1
MEIS1
CDH2
BCL2
CD34
TAL1
DNTT
MME
CD5
CD83
CCND1
TCF3
EBF1
PAX5
CD79A
BTG1
CD19
CD79B
MYBL2
EZH2
HMGB1
HMGB2
BIRC5
CD38
MS4A1
B
Chromatin accessibility
TAL1_chr1:47231645-47232835
CD5_chr11:61102149-61102802
CDH2_chr18:28177958-28178504
MEIS1_chr2:66432902-66435519
CD79B_chr17:63931373-63931667
EBF1_chr5:159099168-159099389
MYBL2_chr20:43666585-43668155
PAX5_chr9:37034185-37035344
DNTT_chr10:96304303-96304541
MS4A1_chr11:60455234-60456087
CD19_chr16:28930614-28932373
CD79B_chr17:63931962-63933283
CD79A_chr19:41876821-41878102
CD38_chr4:15777056-15779532
EZH2_chr7:148883193-148885048
BCL2_chr18:63319534-63321103
HMGB1_chr13:30616967-30618886
BTG1_chr12:92145593-92146594
NOTCH1_chr9:136545144-136547051
TCF3_chr19:1652643-1653384
CD83_chr6:14117594-14118205
HMGB2_chr4:173334011-173335589
BIRC5_chr17:78213977-78214598
CCND1_chr11:69640963-69641388
C
TFBS activity score
Z-score
20
10
0
−10
−20
SPIB
IRF4
SPI1
BCL11A
JUND
FLI1
PAX5
FOXO4
FOXO1
GFI1B
IKZF1
TAL1
GATA1
GFI1
MYBL2
MLXIP
RUNX3
ELK3
ETV6
ETS2
ERG
E2F3
RUNX1
EBF1
TCF4
TCF3
LEF1
MYB
Cell subpopulation
HSC
CLP
pro-B
pre-B
Immature B
Transitional B
Naive CD5+
Naive CD5-
Pattern
No or low expression/accessibility
High expression/accessibility
OCR type
Promoter
Intronic
Distal Intergenic
D
TFs included in each csGRN
TFs included in the csGRN
TFs not included in the csGRN

Figure S7

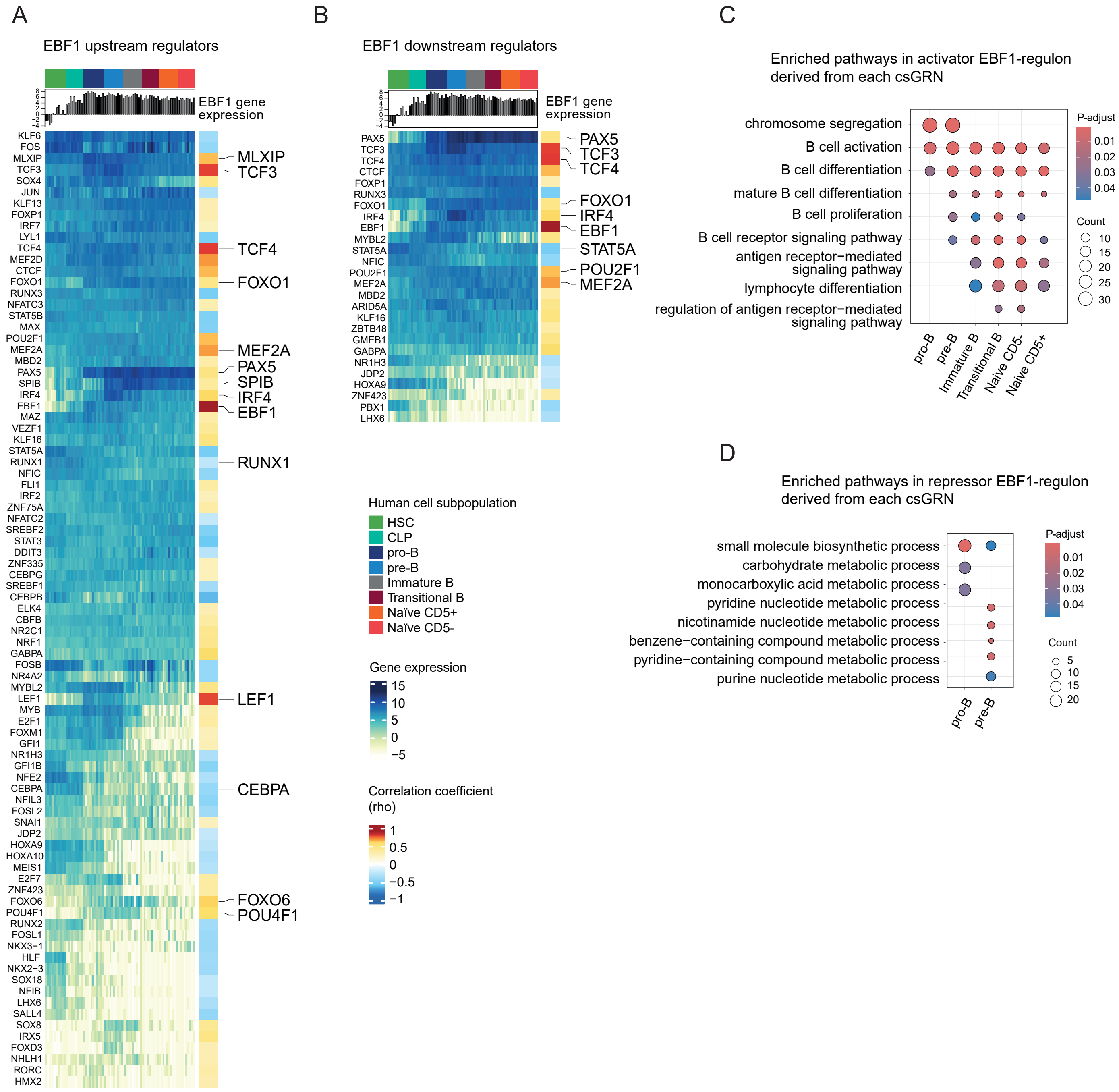
A
EBF1 upstream regulators
EBF1 gene expression
KLF6
FOS
MLXIP
TCF3
SOX4
JUN
KLF13
FOXP1
IRF7
LYL1
TCF4
MEF2D
CTCF
FOXO1
RUNX3
NFATC3
STAT5B
MAX
POU2F1
MEF2A
MBD2
PAX5
SPIB
IRF4
EBF1
MAZ
VEZF1
KLF16
STAT5A
RUNX1
NFIC
FLI1
IRF2
ZNF75A
NFATC2
SREBF2
STAT3
DDIT3
ZNF335
CEBPG
SREBF1
CEBPB
ELK4
CBFB
NR2C1
NRF1
GABPA
FOSB
NR4A2
MYBL2
LEF1
MYB
E2F1
FOXM1
GFI1
NR1H3
GFI1B
NFE2
CEBPA
NFIL3
FOSL2
SNAI1
JDP2
HOXA9
HOXA10
MEIS1
E2F7
ZNF423
FOXO6
POU4F1
RUNX2
FOSL1
NKX3−1
HLF
NKX2−3
SOX18
NFIB
LHX6
SALL4
SOX8
IRX5
FOXD3
NHLH1
RORC
HMX2
MLXIP
TCF3
TCF4
FOXO1
MEF2A
PAX5
SPIB
IRF4
EBF1
RUNX1
LEF1
CEBPA
FOXO6
POU4F1
B
EBF1 downstream regulators
EBF1 gene expression
PAX5
TCF3
TCF4
CTCF
FOXP1
RUNX3
FOXO1
IRF4
EBF1
MYBL2
STAT5A
NFIC
POU2F1
MEF2A
MBD2
ARID5A
KLF16
ZBTB48
GMEB1
GABPA
NR1H3
JDP2
HOXA9
ZNF423
PBX1
LHX6
PAX5
TCF3
TCF4
FOXO1
IRF4
EBF1
STAT5A
POU2F1
MEF2A
Human cell subpopulation
HSC
CLP
pro-B
pre-B
Immature B
Transitional B
Naïve CD5+
Naïve CD5-
Gene expression
15
10
5
0
−5
Correlation coefficient (rho)
1
0.5
0
−0.5
−1
C
Enriched pathways in activator EBF1-regulon derived from each csGRN
chromosome segregation
B cell activation
B cell differentiation
mature B cell differentiation
B cell proliferation
B cell receptor signaling pathway
antigen receptor−mediated signaling pathway
lymphocyte differentiation
regulation of antigen receptor−mediated signaling pathway
pro-B
pre-B
Immature B
Transitional B
Naive CD5-
Naive CD5+
P-adjust
0.01
0.02
0.03
0.04
Count
10
15
20
25
30
D
Enriched pathways in repressor EBF1-regulon derived from each csGRN
small molecule biosynthetic process
carbohydrate metabolic process
monocarboxylic acid metabolic process
pyridine nucleotide metabolic process
nicotinamide nucleotide metabolic process
benzene−containing compound metabolic process
pyridine−containing compound metabolic process
purine nucleotide metabolic process
pro-B
pre-B
P-adjust
0.01
0.02
0.03
0.04
Count
5
10
15
20

Figure S8

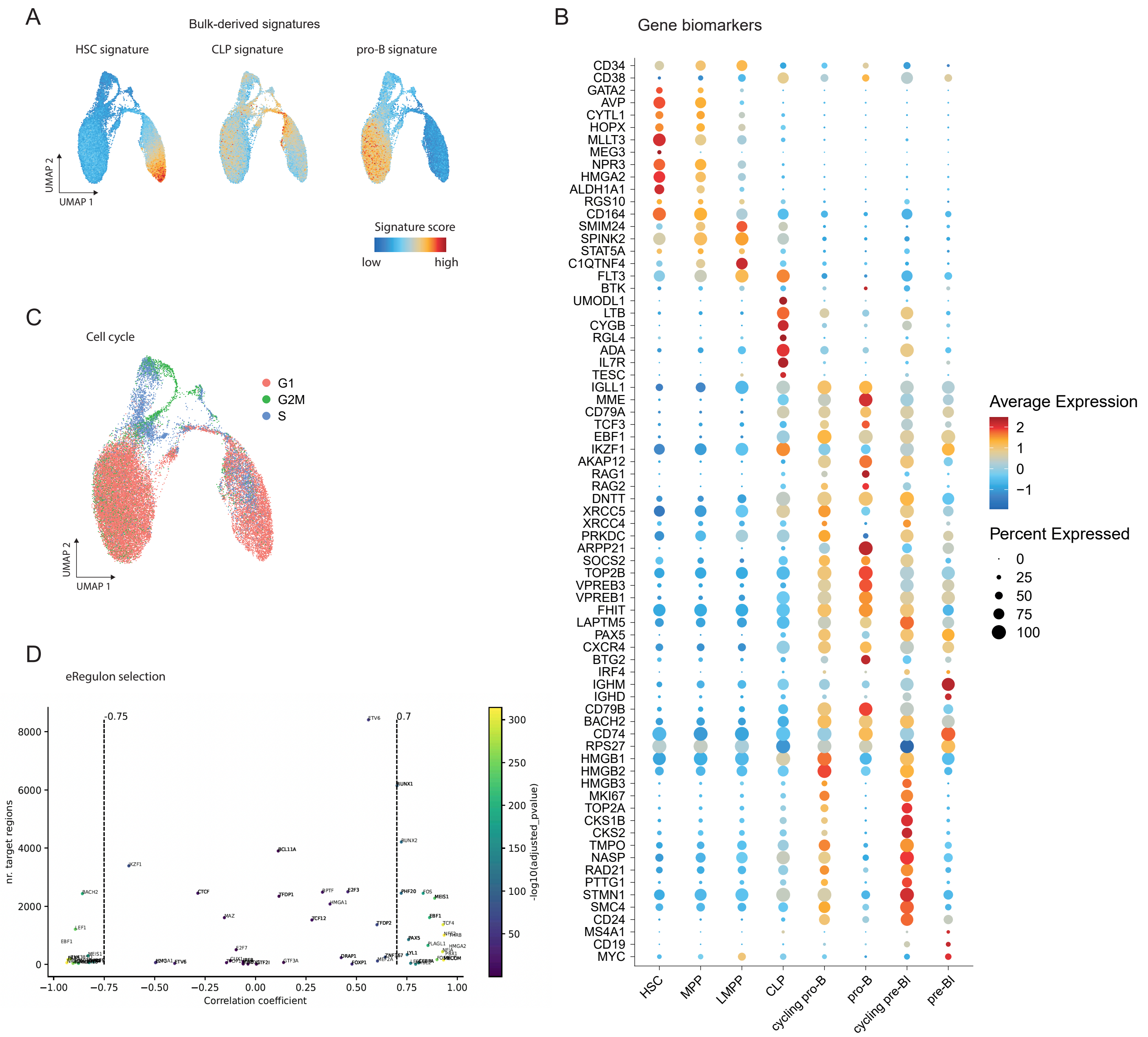
A
Bulk-derived signatures
HSC signature
CLP signature
pro-B signature
UMAP 2
UMAP 1
Signature score
low
high
B
Gene biomarkers
CD34
CD38
GATA2
AVP
CYTL1
HOPX
MLLT3
MEG3
NPR3
HMGA2
ALDH1A1
RGS10
CD164
SMIM24
SPINK2
STAT5A
C1QTNF4
FLT3
BTK
UMODL1
LTB
CYGB
RGL4
ADA
IL7R
TESC
IGLL1
MME
CD79A
TCF3
EBF1
IKZF1
AKAP12
RAG1
RAG2
DNTT
XRCC5
XRCC4
PRKDC
ARPP21
SOCS2
TOP2B
VPREB3
VPREB1
FHIT
LAPTM5
PAX5
CXCR4
BTG2
IRF4
IGHM
IGHD
CD79B
BACH2
CD74
RPS27
HMGB1
HMGB2
HMGB3
MKI67
TOP2A
CKS1B
CKS2
TMPO
NASP
RAD21
PTTG1
STMN1
SMC4
CD24
MS4A1
CD19
MYC
HSC
MPP
LMPP
CLP
cycling pro-B
pro-B
cycling pre-BI
pre-BI
Average Expression
2
1
0
−1
Percent Expressed
0
25
50
75
100
C
Cell cycle
G1
G2M
S
UMAP 2
UMAP 1
D
eRegulon selection
-0.75
0.7
8000
6000
4000
2000
0
nr. target regions
−1.00
−0.75
−0.50
−0.25
0.00
0.25
0.50
0.75
1.00
Correlation coefficient
300
250
200
150
100
50
-log10(adjusted_pvalue)

### Figure S9

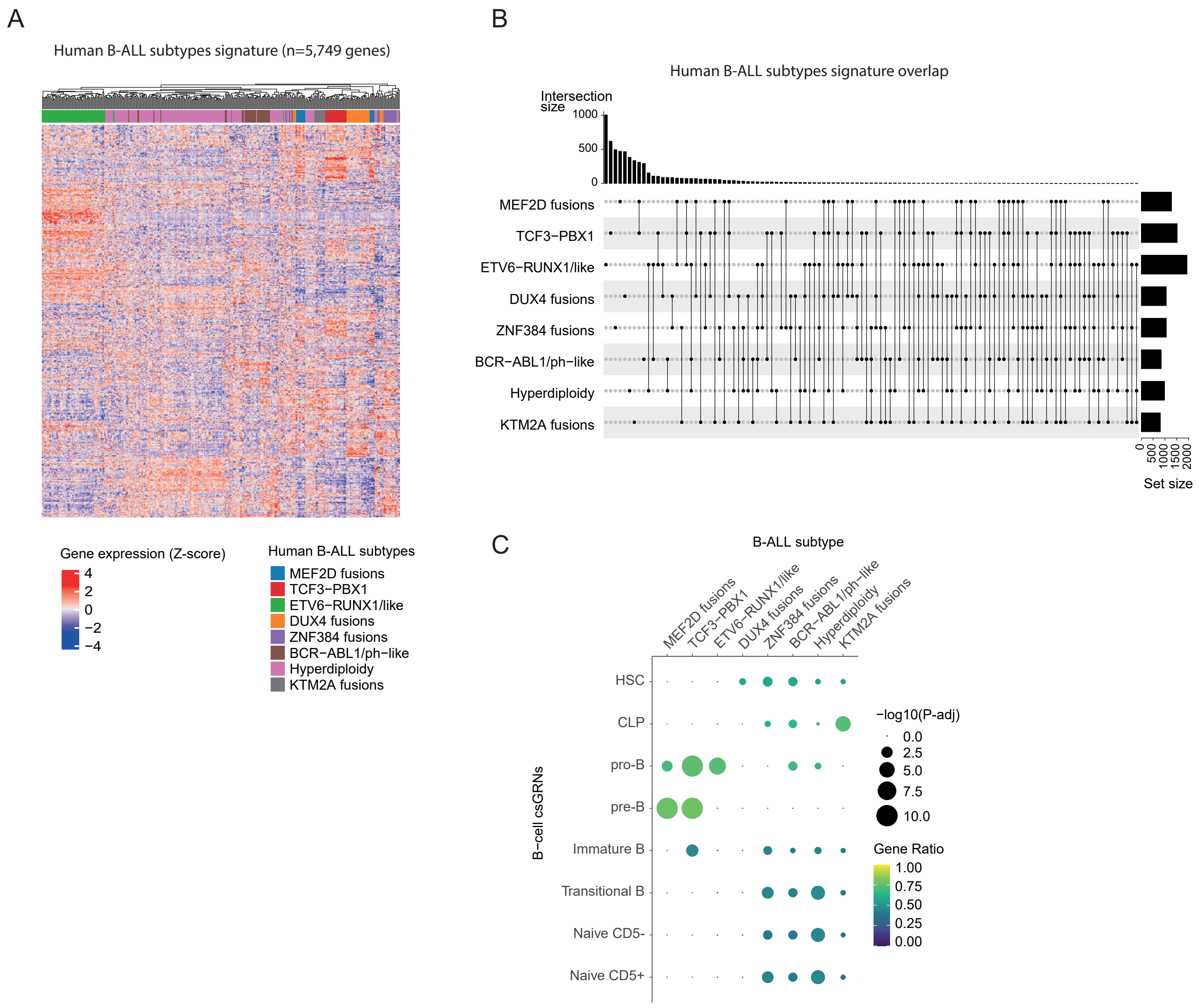

A
Human B-ALL subtypes signature (n=5,749 genes)
Gene expression (Z-score)
4
2
0
−2
−4
Human B-ALL subtypes
MEF2D fusions
TCF3−PBX1
ETV6−RUNX1/like
DUX4 fusions
ZNF384 fusions
BCR−ABL1/ph−like
Hyperdiploidy
KTM2A fusions
B
Human B-ALL subtypes signature overlap
Intersection size
1000
500
0
MEF2D fusions
TCF3−PBX1
ETV6−RUNX1/like
DUX4 fusions
ZNF384 fusions
BCR−ABL1/ph−like
Hyperdiploidy
KTM2A fusions
0
500
1000
1500
2000
Set size
C
B-ALL subtype
MEF2D fusions
TCF3−PBX1
ETV6−RUNX1/like
DUX4 fusions
ZNF384 fusions
BCR−ABL1/ph−like
Hyperdiploidy
KTM2A fusions
B−cell csGRNs
HSC
CLP
pro-B
pre-B
Immature B
Transitional B
Naive CD5-
Naive CD5+
−log10(P-adj)
0.0
2.5
5.0
7.5
10.0
Gene Ratio
1.00
0.75
0.50
0.25
0.00

Figure S10

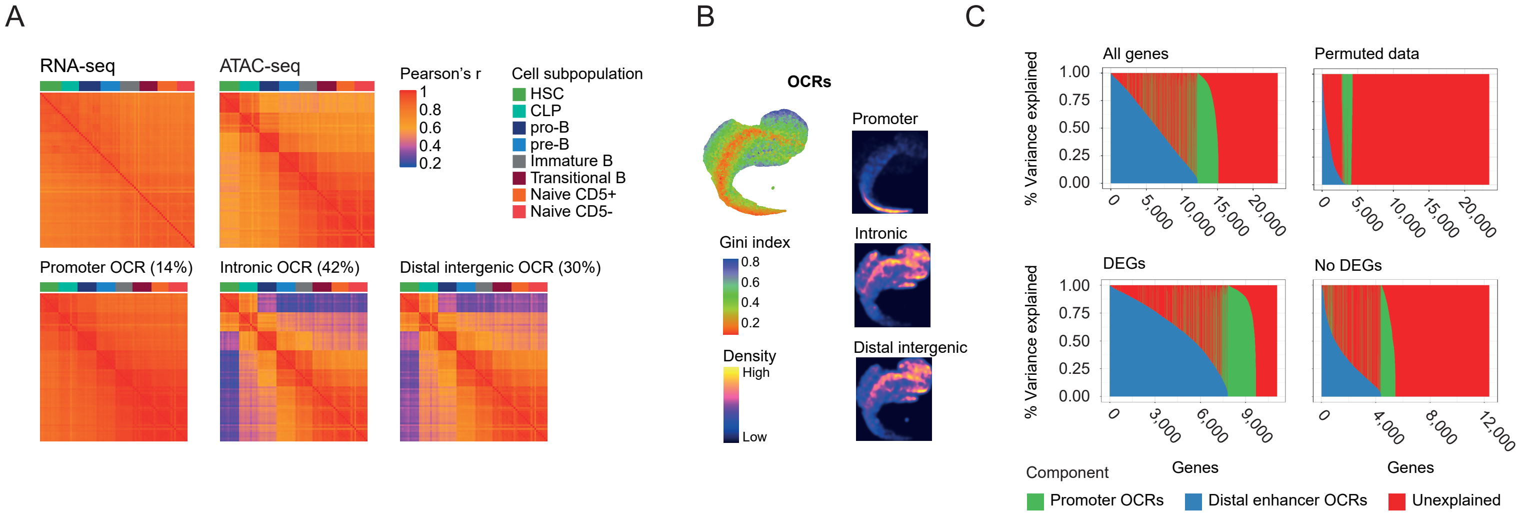
A
RNA-seq
ATAC-seq
Pearson's r
1
0.8
0.6
0.4
0.2
Cell subpopulation
HSC
CLP
pro-B
pre-B
Immature B
Transitional B
Naive CD5+
Naive CD5-
Promoter OCR (14%)
Intronic OCR (42%)
Distal intergenic OCR (30%)
B
OCRs
Promoter
Intronic
Distal intergenic
Gini index
0.8
0.6
0.4
0.2
Density
High
Low
C
All genes
Permuted data
DEGs
No DEGs
% Variance explained
1.00
0.75
0.50
0.25
0.00
0
5,000
10,000
15,000
20,000
3,000
6,000
9,000
4,000
8,000
12,000
Genes
Component
Promoter OCRs
Distal enhancer OCRs
Unexplained

Figure S11

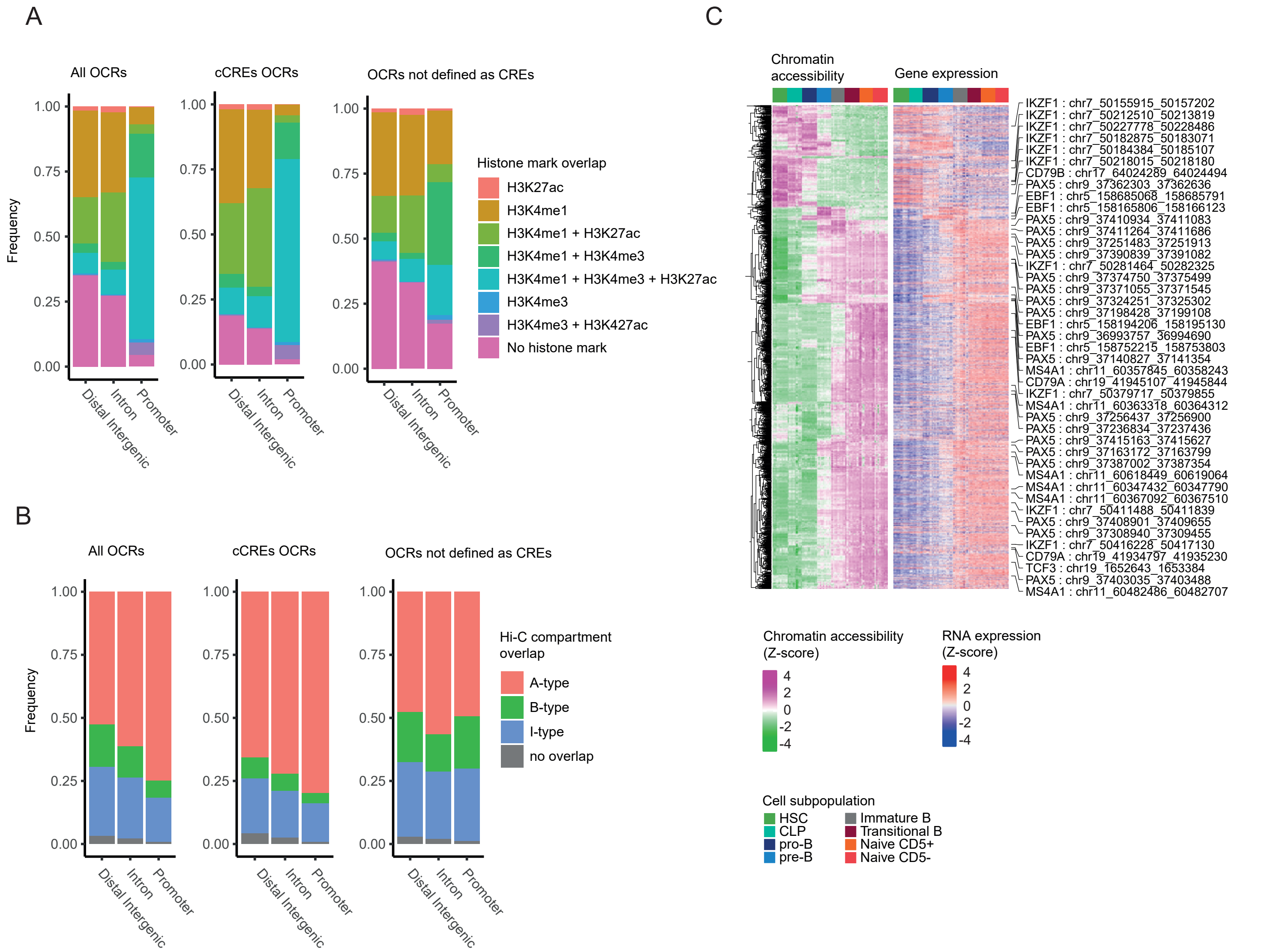

A
All OCRs
cCREs OCRs
OCRs not defined as CREs
Frequency
1.00
0.75
0.50
0.25
0.00
Distal Intergenic
Intron
Promoter
Histone mark overlap
H3K27ac
H3K4me1
H3K4me1 + H3K27ac
H3K4me1 + H3K4me3
H3K4me1 + H3K4me3 + H3K27ac
H3K4me3
H3K4me3 + H3K427ac
No histone mark
B
All OCRs
cCREs OCRs
OCRs not defined as CREs
Frequency
Hi-C compartment overlap
A-type
B-type
I-type
no overlap
C
Chromatin accessibility
Gene expression
IKZF1 : chr7_50155915_50157202
IKZF1 : chr7_50212510_50213819
IKZF1 : chr7_50227778_50228486
IKZF1 : chr7_50182875_50183071
IKZF1 : chr7_50184384_50185107
IKZF1 : chr7_50218015_50218180
CD79B : chr17_64024289_64024494
PAX5 : chr9_37362303_37362636
EBF1 : chr5_158685068_158685791
EBF1 : chr5_158165806_158166123
PAX5 : chr9_37410934_37411083
PAX5 : chr9_37411264_37411686
PAX5 : chr9_37251483_37251913
PAX5 : chr9_37390839_37391082
IKZF1 : chr7_50281464_50282325
PAX5 : chr9_37374750_37375499
PAX5 : chr9_37371055_37371545
PAX5 : chr9_37324251_37325302
PAX5 : chr9_37198428_37199108
EBF1 : chr5_158194206_158195130
PAX5 : chr9_36993757_36994690
EBF1 : chr5_158752215_158753803
PAX5 : chr9_37140827_37141354
MS4A1 : chr11_60357845_60358243
CD79A : chr19_41945107_41945844
IKZF1 : chr7_50379717_50379855
MS4A1 : chr11_60363318_60364312
PAX5 : chr9_37256437_37256900
PAX5 : chr9_37236834_37237436
PAX5 : chr9_37415163_37415627
PAX5 : chr9_37163172_37163799
PAX5 : chr9_37387002_37387354
MS4A1 : chr11_60618449_60619064
MS4A1 : chr11_60347432_60347790
MS4A1 : chr11_60367092_60367510
IKZF1 : chr7_50411488_50411839
PAX5 : chr9_37408901_37409655
PAX5 : chr9_37308940_37309455
IKZF1 : chr7_50416228_50417130
CD79A : chr19_41934797_41935230
TCF3 : chr19_1652643_1653384
PAX5 : chr9_37403035_37403488
MS4A1 : chr11_60482486_60482707
Chromatin accessibility (Z-score)
4
2
0
-2
-4
RNA expression (Z-score)
4
2
0
-2
-4
Cell subpopulation
HSC
CLP
pro-B
pre-B
Immature B
Transitional B
Naive CD5+
Naive CD5-
